# Spatial Transcriptomics Reveals Expression Gradients in Developing Wheat Inflorescences at Cellular Resolution

**DOI:** 10.1101/2024.12.19.629411

**Authors:** Katie A. Long, Ashleigh Lister, Maximillian R. W. Jones, Nikolai M. Adamski, Rob E. Ellis, Carole Chedid, Sophie J. Carpenter, Xuemei Liu, Anna E. Backhaus, Andrew Goldson, Vanda Knitlhoffer, Yuanrong Pei, Martin Vickers, Burkhard Steuernagel, Gemy G. Kaithakottil, Jun Xiao, Wilfried Haerty, Iain C. Macaulay, Cristobal Uauy

## Abstract

The diversity of plant inflorescence architectures is specified by gene expression patterns. In wheat (*Triticum aestivum*), the lanceolate-shaped inflorescence (spike) is defined by rudimentary spikelets at the base which initiate first but subsequently lag in development compared with central spikelets. While previous studies identified gene expression differences between central and basal inflorescence sections, the spatio-temporal dynamics and gradients along the apical-basal axis remain poorly resolved due to bulk tissue-level techniques. Here, using spatial transcriptomics, we profiled 200 genes across four stages of wheat inflorescence development to cellular resolution. Cell segmentation and unsupervised clustering identified 18 expression domains and their enriched genes, revealing dynamic spatio-temporal organisation along the apical-basal axis of the inflorescence. Along this axis, we uncovered distinct and spatially coordinated gene expression gradients patterning meristems prior to the visible delay in basal spikelet development. This study demonstrates the potential for spatial transcriptomics time-series to advance plant developmental biology.

## Introduction

Nature provides a stunning diversity of vegetative and flowering structures contributing to plant fitness. Understanding the process of tissue patterning requires characterizing regulatory genes and their spatial expression, as cell fate depends on positional cues within developing tissues^1–4^. Grass morphology is patterned through phytomers, a basic unit consisting of a node, internode, leaf, and axillary meristem (AM)^5,6^. During vegetative growth, the shoot apical meristem (SAM) initiates phytomers, with lateral leaf primordia outgrowing while AMs remain dormant^7^. This developmental trajectory shifts as the SAM transitions to the inflorescence meristem. Phytomer initiation continues, however leaf (bract) outgrowth is suppressed, and AMs transition to spikelet meristems (SM) to pattern reproductive growth^8,9^.

Bread wheat (*Triticum aestivum*) forms an unbranched spike-type inflorescence where pairs of AMs and suppressed bracts are initiated sequentially forming a gradient of meristem ages along the inflorescence, with basal meristems being the oldest ^10,11^. The timing of transition of each AM (spikelet ridges (SRs) in wheat) into SMs, however, does not align with their developmental progression. Central SRs along the apical-basal axis are the first to initiate spikelet development, while basal SRs, despite having more time to develop, lag behind^11^. This delay persists throughout inflorescence patterning, resulting in basal spikelets forming immature floral structures that fail to produce grains^12,13^. This highlights the composite nature of the developing wheat spike, where meristems of different ages and developmental stages coexist within an ∼1000 µm inflorescence, ultimately influencing its final structure.

Low-input RNA-seq of apical, central and basal sections of micro-dissected wheat spikes revealed large differences in gene expression profiles among them^13^. For example, the MADS-box transcription factor *VEGETATIVE TO REPRODUCTIVE TRANSITION 2 (VRT2)* exhibited its highest expression in basal sections, with decreasing levels toward the apex, in a proposed gradient along the inflorescence. Increased expression of *VRT2* led to a subtle delay in basal spikelet development and elongated organs within the spikelet (glumes and lemmas)^13,14^. While informative, the semi-spatial resolution of microdissections leaves precise expression patterns and putative gradients undefined. Given the high levels of differential expression observed across the spike in this experiment, we hypothesized that other genetic factors contribute to apical-basal axis patterning that warrant further investigation.

Here, we employ an adapted Multiplexed Error Robust Fluorescence *In Situ* Hybridization (MERFISH)^15,16^ protocol to map the spatial expression of 200 genes along the apical-basal axis of the wheat spike to cellular resolution. Cell segmentation and unsupervised clustering across four timepoints identified 18 expression domains and their enriched gene markers, offering detailed insights into gene expression at tissue and cellular levels^17–20^. We uncovered detailed spatial and temporal organization patterns in the developing wheat spike, including coordinated transcriptional gradients that distinguish and define leaf and spikelet ridges along the apical-basal axis, prior to central meristem outgrowth. To support the broader research community, we developed an open-access WebAtlas interface^21^ (www.wheat-spatial.com), enabling visualization of all measured genes and expression domains. This work highlights the power of spatial transcriptomics time-series to investigate gene expression patterns to single cell resolution while retaining the tissue morphology and spatial context of each cell *in planta*.

## Results

### MERFISH of wheat inflorescence resolves gene expression to cellular resolution across four developmental timepoints

To study transcriptional gradients in wheat spikes over time, we selected genes of interest for the MERSCOPE spatial transcriptomics platform. To define these genes we first examined a micro-dissection RNA-seq dataset^13^, however, given its limited developmental range and high variability, we conducted a more extensive analysis across spike development. We generated RNA-seq from central and basal spike sections across five development stages (Extended Data Fig. 1a,b; Supplementary Table 1)^22^; Early and Late double ridge (EDR, LDR; Waddington stage W2, W2.5; respectively), Lemma Primordia (LP; W3.25), Terminal Spikelet (TS; W4), and Carpel Extension (CE; W5). Individual samples expressed, on average, 49,387 high-confidence genes, with 55,346 unique genes expressed across all samples. We identified 12,384 genes with significant differential expression between central and basal sections over time (p_adj_ < 0.001; Extended Data Fig. 1c; Supplementary Table 2), consistent with distinct developmental pathways along the apical-basal axis.

We selected 116 genes from our differential expression dataset, supplemented with 73 additional genes including known inflorescence development genes. The final panel included 189 inflorescence related genes, 100 grain development genes for a separate project, and eleven house-keeping genes (Supplementary Table 3). After designing and synthesising the 300-probe set, we performed the Vizgen MERSCOPE workflow (see Methods). Developing wheat spikes were dissected at four stages (W2.5, W3.25, W4, W5), embedded in optimal cutting temperature (OCT) compound, flash frozen, and cryo-sectioned (Extended Data Fig. 1d). Each OCT block included 5–36 spikes (depending on stage) from two near-isogenic lines: one carrying the wildtype *VRT-A2a* allele (*P1^WT^*) and the other the misexpression *VRT-A2b* allele from *T. turgidum* ssp. *polonicum*^14^ (*P1^POL^*; Extended Data Fig. 1e; Supplementary Fig. 1). We hybridised the probes to the samples and imaged across five experimental runs.

For downstream analysis, we selected two representative samples at each timepoint (one per genotype) for a total of eight samples. Cell segmentation^17^ and transcript assignment^23^ (Fig. 1a-e) yielded a cell-by-gene matrix detailing transcript counts per cell for each gene (Supplementary Table 4). After filtering for low-quality cells (see Methods), we assessed sample quality using ‘total transcript counts’ and ‘gene counts per cell’ metrics. Across all samples, the total transcript counts per cell averaged between 77.9 to 152.4 counts, while average gene counts per cell ranged from 29.6 to 42.1 (Supplementary Fig. 2a,b, Supplementary Table 5). We observed gene expression patterns at cellular resolution, including transcripts localized to a single cell layer of the epidermis (Fig. 1f) and clear separation of floral tissues by the profiles of floral homeotic regulators (e.g. *AGAMOUS-LIKE6* (*AGL6*), *PISTILLATA1* (*PI1*); Fig. 1g). These observations are consistent with MERFISH providing spatial expression data at cellular resolution.

**Figure 1.**
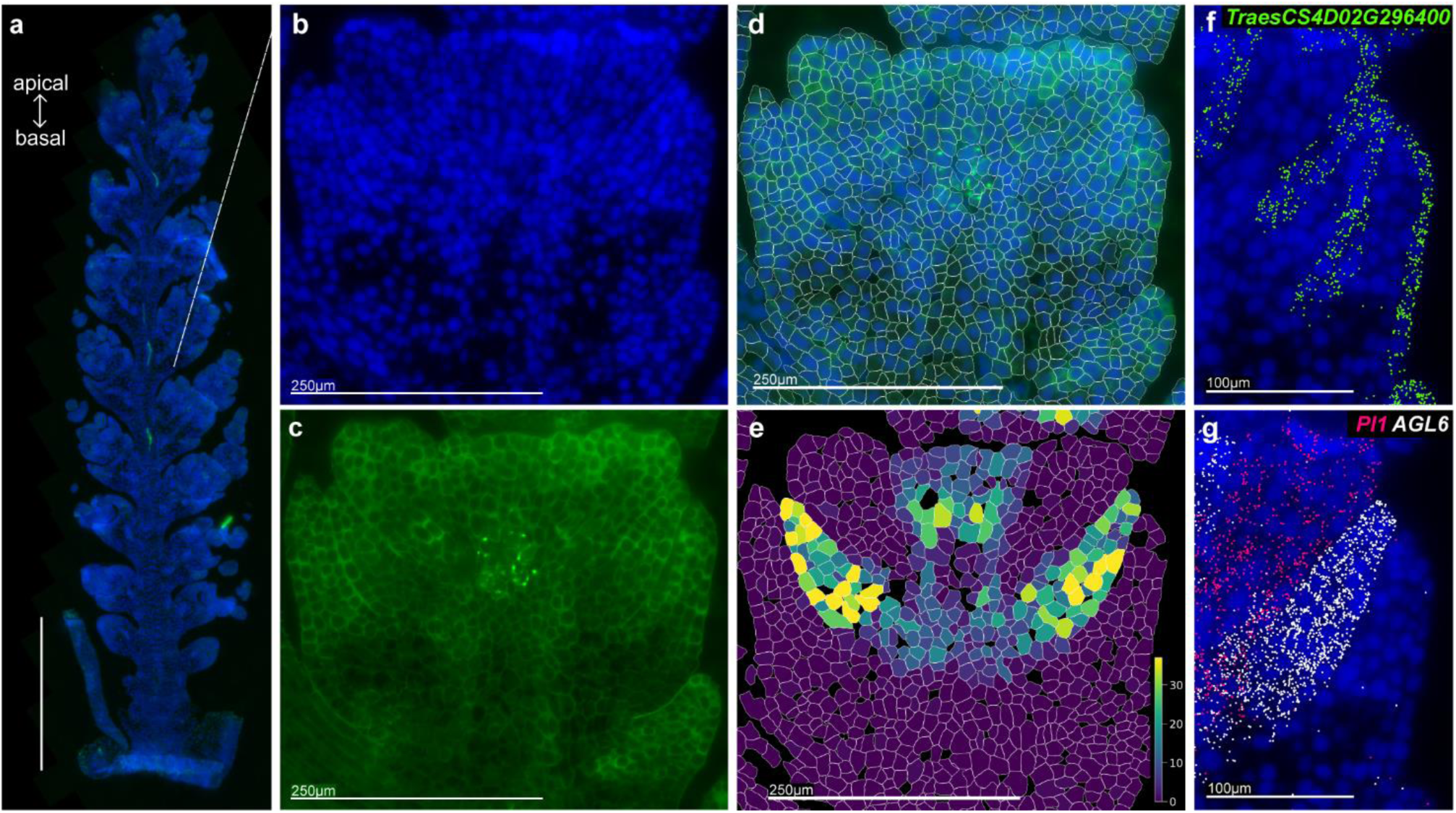
Development and implementation of a 200-gene MERFISH panel in wheat inflorescence tissue. **a**, Cryosection of W5 spike showing DAPI stain (blue) and polyT stain (green). Inset: higher magnification of floral tissues with individual staining of DAPI (**b**) and PolyT (**c**). **d,** Cellular segmentation with cellpose2, filtered for high quality cells. **e**, Heatmap displaying ‘transcript counts per cell’ for *AGL6*, with assignment conducted using the Vizgen Post-Processing Tool. **f**, *TRAESCS4D02G296400* transcripts (green) localized in the first cell layer of floral tissues. **g**, Tissue-specific expression patterns of *AGL6* (white) and *PI1* (pink) with distinct, non-overlapping spatial localization towards tips of palea and stamen, respectively.

### MERFISH yields low off-target rates and consistent expression patterns with *in situ* hybridisations

We next implemented quality control measures to evaluate the performance of MERFISH in plants. We detected minimal off-target hybridisation based on 15 blank controls which represented <0.3% of total counts per cell (range: 0.24%–0.35%; Supplementary Table 6). As expected, probes were non-homoeolog specific, an important consideration for polyploids (Supplementary Note 1; Extended Data Fig. 2). To further validate our results, we generated *in silico* sections equivalent to physical microdissections used in RNA-seq, yielding an average Spearman’s correlation coefficient of 0.66 (range 0.60-0.73; Supplementary Fig. 2c, Supplementary Table 7), supporting consistency between approaches. Additionally, MERFISH data was consistent with published *in situ* hybridizations in cereals (Supplementary Table 8, Supplementary Fig. 3). These results demonstrate that our MERFISH data exhibits minimal off-target activity, is non-homoeolog specific, and is consistent with established gene expression patterns, confirming the technique’s robustness in plant tissues.

### Unsupervised clustering defines 18 expression domains traced through developmental time

We integrated eight samples and applied unsupervised clustering^19,18^ which identified 18 distinct expression domains (Fig. 2a). These were visualized as spatially resolved maps across developmental stages and genotypes (Fig. 2b-e; Extended Data Fig. 3). By concatenating all time points before clustering, we traced the spatio-temporal dynamics of the domains. Notably, six domains were primarily detected at stages W4 and W5 (Fig. 2f; Supplementary Table 9), which we identified as floral tissues (Fig. 2a-e) based on their spatial distribution and organization. We hypothesised that concatenating samples across time would reveal cells in their earliest stages of differentiation. For example, in W3.25 spikes, we identified only three expression domain 15 (ED15) cells (Fig. 2g), a predominant domain in mature spikes and which later localises to paleae in W5 samples (Fig. 2h). At W3.25, these three ED15 cells were positioned just above the lemma primordia (ED1, ED2), a spatial arrangement consistent across stages. ED15 cells expressed *AGL6* (Extended Data Fig. 4), suggesting an early role for *AGL6* in palea identity consistent with wheat *agl6* mutants where paleae transform into lemma-like organs^24^. This indicates that ED15 at W3.25 represents the earliest palea progenitor cells. The presence of ED15 cells on the seventh spikelet ridge again highlights developmental differences between central and basal spikelets and the composite nature of meristems along the spike.

**Figure 2.**
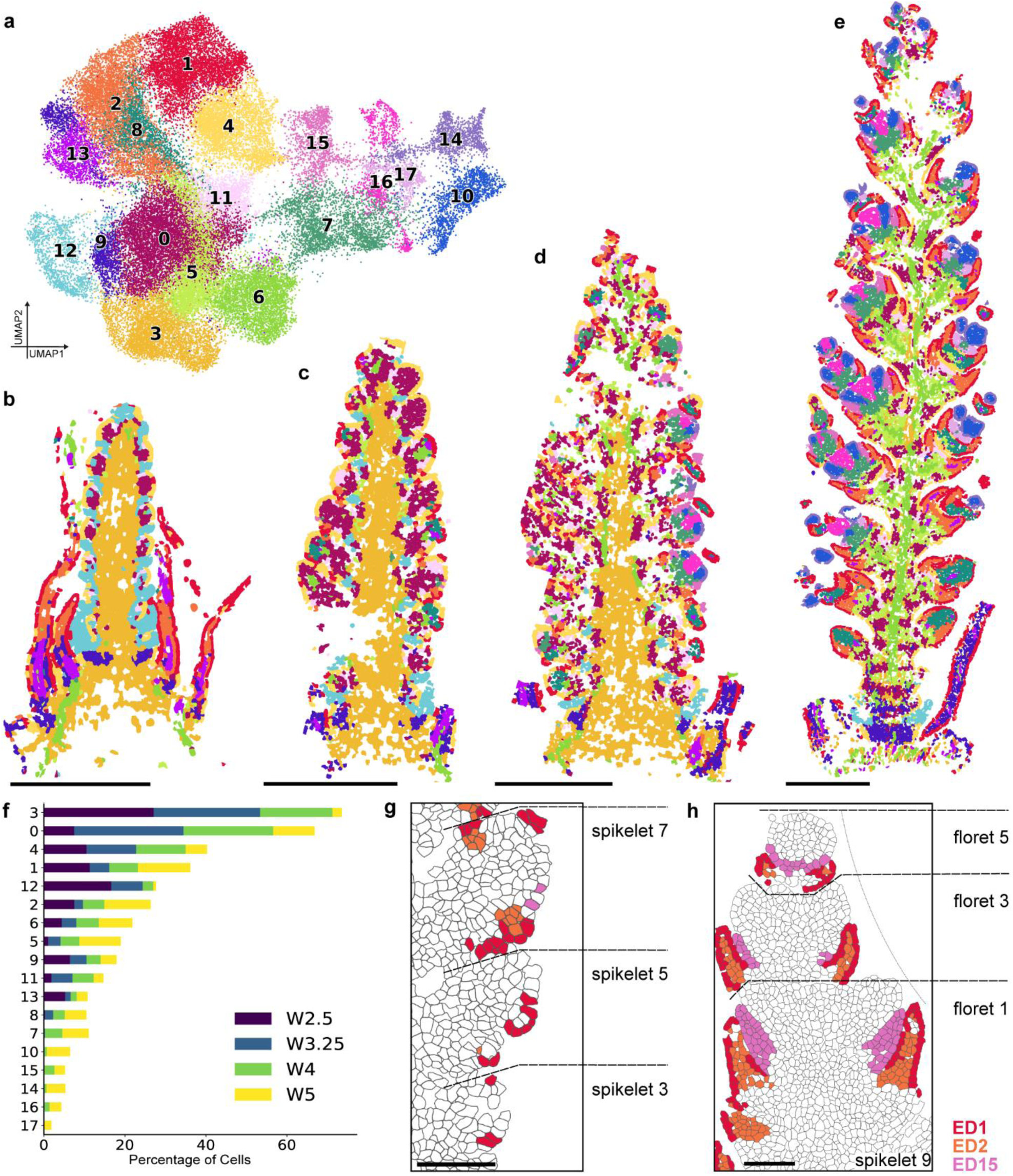
Unsupervised clustering identifies 18 expression domains mapped over four developmental stages. **a**, UMAP projection of cells from eight samples, and expression domain assignment. **b-e**, Spatial maps of Leiden clustering across time points W2.5 (**b**), W3.25 (**c**), W4 (**d**), and W5 (**e**) using Squidpy (v1.4.1), Scanpy (v 1.10.0), and Scanorama (v1.7.4) scale bar = 500µm. f, Proportion of cells in each expression domain per sample, calculated as the percentage of total cells assigned to each cluster. **g-h**, Spatial plots of cell segmentation and assigned expression domains 1, 2, & 15 in (**g**) W3.25 spikelets and (**h**) W5 florets highlight the tracing of cell groups across developmental time, with two ED15 cells in W2.5 found above glumes & lemmas (ED1&2) tracing to palea in W5 florets. Scale bar = 100µm.

### Gene Expression Analysis defines identity of Expression Domains

Expression domains require biological context for tissue-specific annotation. We annotated the 18 domains by combining anatomical features, such as ED15’s association with palea, with domain-enriched genes (Methods; Supplementary Table 10-11). These annotations remained consistent across time and genotypes. For instance, at W2.5, ED12 is enriched for *TraesCS1A02G418200* (Fig. 3a-b), the wheat ortholog of maize *TASSEL SHEATH1* (*TSH1*) which suppresses bract outgrowth in cereal inflorescences^8,25–28^. Furthermore, at W2.5 ED12 cells and *TSH1* expression form a repeating pattern along the spike, corresponding to the suppressed leaf ridge (LR). By W3.25, LRs are no longer visible in the central and apical spike sections, yet *TSH1* persists in the few ED12 cells at the base of each spikelet ridge (SR), indicating active LR suppression during early floral organ differentiation. Furthermore, we validated the ABCDE model of floral development in wheat (Supplementary Note 2; Extended Data Fig. 5; Supplementary Table 12).

**Figure 3.**
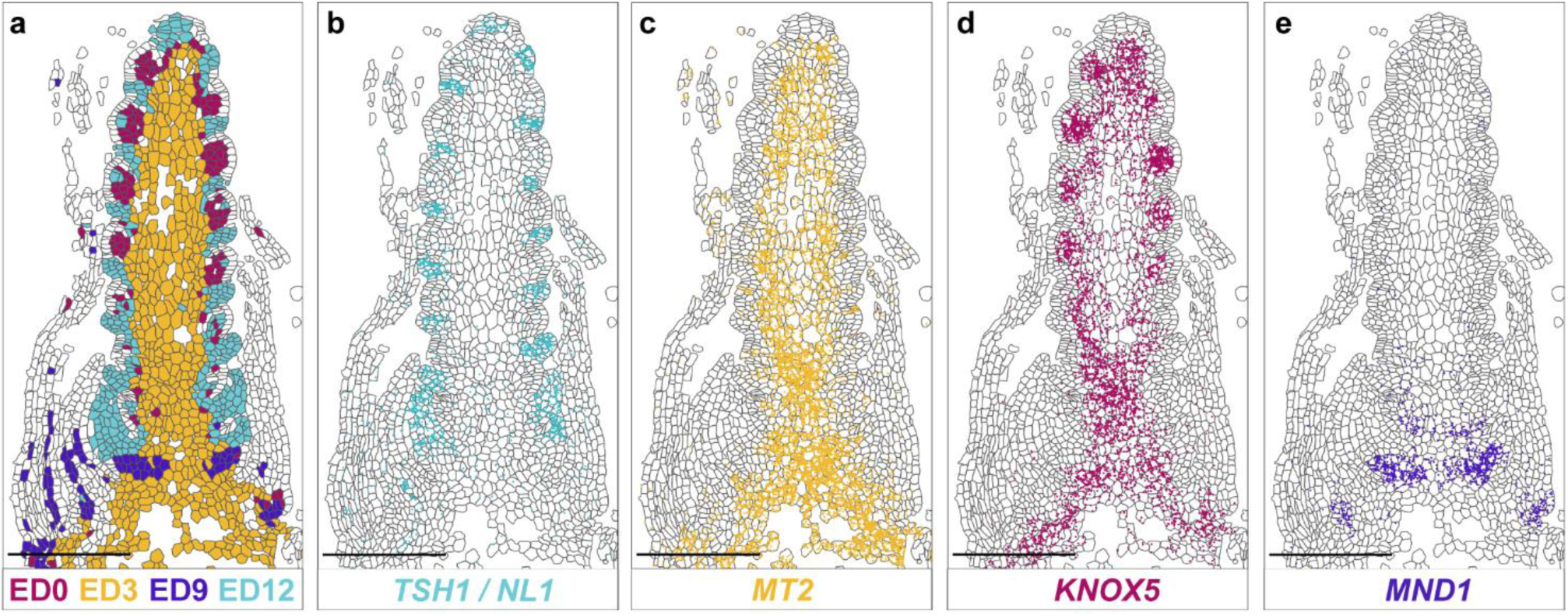
Gene expression analysis defines identity and enriched genes in Expression Domains. **a**, Spatial map of four Expression Domains in W2.5 spikes, highlighting domains enriched with transcripts of (**b**) *TASSEL SHEATH1 (TSH1)*/ *NECK LEAF 1 (NL1)*; (**c**) *METALLOTHIONEIN 2 (MT2)*; (**d**) *KNOTTED1-LIKE HOMEOBOX 5 (KNOX5)*, and (**e**) *MANY NODED DWARF 1 (MND1)*. Scale bar = 250µm.

Some genes uniquely mark single domains, such as *METALLOTHIONEIN 2* (*MT2*), which identifies early vasculature in ED3 (Fig. 3a,c). Others are enriched across multiple domains, such as *KNOTTED1-LIKE HOMEOBOX 5* (*KNOX5*) in ED3 (developing rachis) and ED0 (meristem cells; Fig. 3a,d). *KNOX5* exclusion from the L1 layer is consistent with observations of its maize ortholog *KNOTTED1*^29^. Certain domains encompass multiple tissues: ED9 marks both developing young leaves and suppressed AM below the inflorescence *per se,* with *MANY NODED DWARF 1* (*MND1*) marking suppressed AMs only (Fig. 3a,e). We hypothesize that the spike-focused probe panel lacked sufficient genes to distinguish these tissues as distinct ED. Despite the limitations of the 200-gene panel, these findings demonstrate how expression domains capture complex gene expression profiles, revealing distinct developmental identities within the spike.

### Spatial analysis of *VRT2* and *SEP1-4* gradients

We previously identified differences in gene expression between spike sections using microdissection. *VRT2* was most highly expressed in basal sections, with lower expression in the apex, whereas *SEPALLATA* (*SEP*) MADS-box transcription factors displayed the opposite pattern^13^. To quantify these profiles at W4, we computationally dissected spikes into 25 transverse bins, revealing opposing expression gradients along the apical-basal axis (Fig. 4a,b). Expression domain analysis revealed spatial segregation of *VRT2* and *SEP1-4*. *VRT2* was primarily expressed in ED3 developing rachis cells with 32.2% of ED3 cells expressing *VRT2* and showed minimal expression in spikelet tissues such as glumes/lemmas (ED2, 3.3%). In contrast, *SEP1-4* was largely absent from ED3 (1.4%) but enriched in spikelet tissues including glumes/lemmas (ED2, 31.1%; Fig. 4c; Supplementary Table 13). Across the spike, only 0.7% of cells co-expressed *VRT2* and *SEP1-4*.

**Figure 4.**
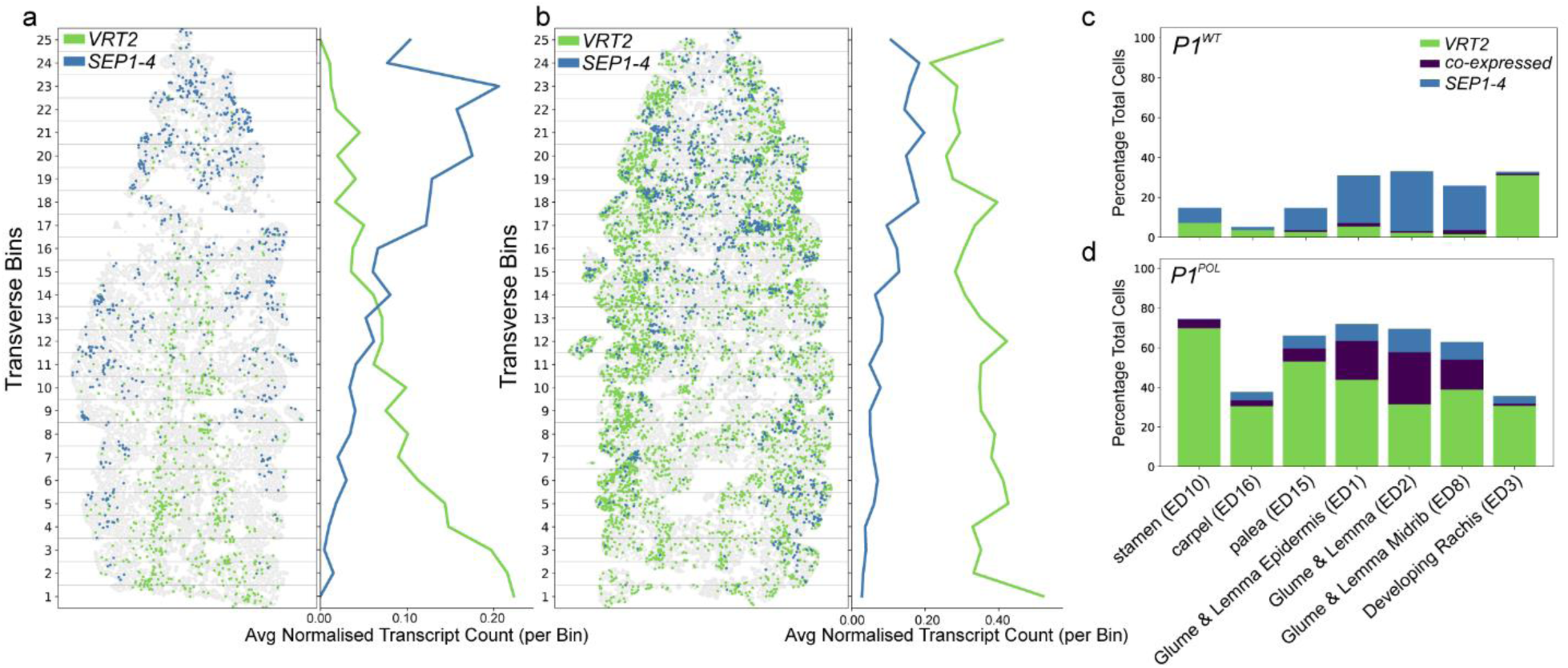
Opposing spatial gradients of *VRT2* and *SEP1-4* along the apical-basal axis disrupted by ectopic expression in *P1^POL^*. **a**, Spatial plot of *VRT2* and *SEP1-4* expression in *P1^WT^* W4 spikes, divided into 25 transverse bins along the apical-basal axis, with average normalized expression counts of *VRT2* and *SEP1-4* per transverse bin, normalised with sc.pp.normalize_total and sc.pp.log1p functions (Scanpy v1.10.0). **b**, Spatial plot of *VRT2* and *SEP1-4* expression in *P1^POL^* W4 spikes, divided into 25 transverse bins along the apical-basal axis. with average normalized expression counts of *VRT2* and *SEP1-4* per transverse bin, normalised with sc.pp.normalize_total and sc.pp.log1p functions (Scanpy v1.10.0). Note difference in scale with (**a**). **c-d**, Proportion of cells expressing either *VRT2, SEP1-4,* or co-expressing both. Calculated as the percentage of total cells per cluster type in (**c**) *P1^WT^* and (**d**) *P1^POL^* isogenic lines.

Next, we analysed the spatial profiles of *VRT2* and *SEP1-4* in *P1^POL^*. MERFISH revealed ectopic *VRT2* expression and disruption of its gradient, whereas the *SEP1-4* gradient remained intact (Fig. 4b). This led to increased co-localisation of *VRT2* and *SEP1-4*, with 8.2% of cells co-expressing both transcripts along the spike. Co-expression was most pronounced in tissues exhibiting the strongest phenotypic effects in *P1^POL^*, glumes and lemmas, where 26.3% of ED2 cells co-expressed both genes (compared to 1.1% in *P1^WT^*; Fig. 4d; Supplementary Table 13). These results support the ‘protein competition’ model proposed by Li et al^30^, where VRT2 interferes with SEP-FRUITFULL2 protein interactions essential for normal spikelet development. Overall, these findings demonstrate the precision of MERFISH in detecting gene expression gradients and uncovering tissue-specific co-expression patterns in developmental mutants.

### Transcriptional states differentiate spikelet meristems and suppressed leaf ridges along the spike

The chronological initiation of SRs does not coincide with their developmental progression. Basal SRs, though first to initiate, lag behind in development compared to central SRs. By the Glume Primordium stage (W3), central SRs display visible outgrowth, while basal SRs remain less developed^11^. We hypothesized that additional gene expression gradients, beyond *VRT2* and *SEP1-4*, may influence these differences before W3. To explore this, we analysed transcriptional and ED patterns in W2.5 spikes. At this stage, the spike has a relatively simple ED composition, with four domains accounting for 94.8% of the inflorescence. By contrast, by W3.25, eight domains account for a comparable proportion (94.1%). The W2.5 SR comprises four domains: the L1 layer (ED4), meristematic cells in layers L2/L3 (ED0, ED12), and boundary cells (ED11) marking the adaxial boundary (Supplementary Table 14). While all SRs exhibit similar L1 (ED4) and boundary (ED11) patterns, basal SRs lack distinct ED0 regions and the expression of *KNOX5* typical of central SRs (Fig. 5a,b). Additionally, basal LR are larger, averaging 32.5 ± 15.8 cells per section (LR1-4), compared to 12.5 ± 1.5 cells in central LRs (LR8-11; Supplementary Table 14). These findings suggest developmental gradients between basal and central SR during or before W2.5.

**Figure 5.**
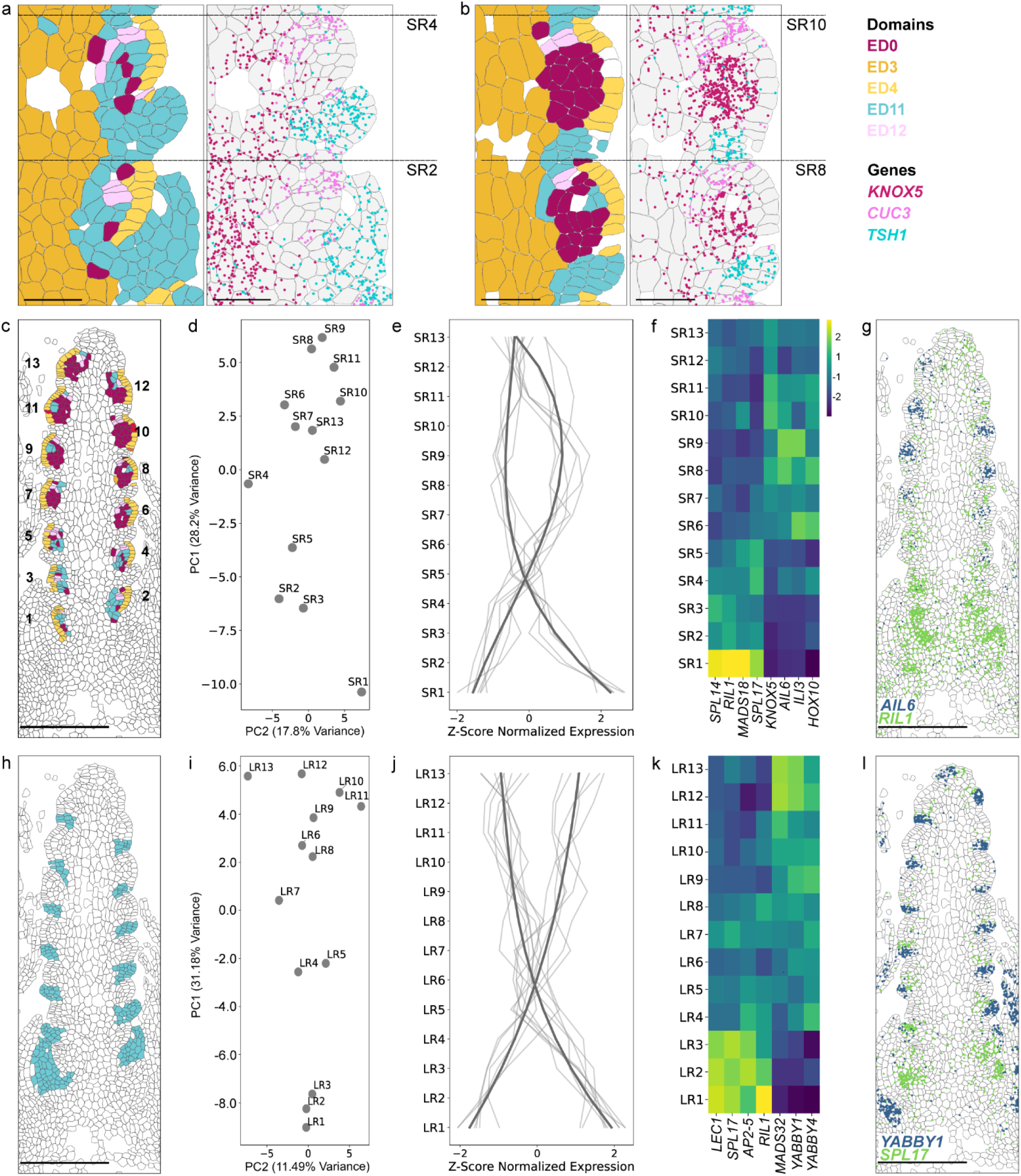
Coordinated and distinct transcriptional gradients define leaf ridges and spikelet ridges across the apical-basal axis at double ridge stage. **a-b**, Basal (2,4) and central (8,10) spikelet ridges (SRs) differ in expression domain assignment and marker gene expression. *TSH1* and *CUC3* mark the suppressed leaf ridge and the adaxial boundary of the spikelet meristem, respectively. *KNOX5* in layers L2/L3 highlights transcriptional differences between SRs along the apical-basal axis. **c**, Expression domains annotate and group SRs from 1 (basal) to 13 (apical). **d**, Principal component (PC) analysis of averaged transcripts per SR group. **e**, Normalized gene expression (Z-score) of genes highly correlated with PC1 (correlation > 0.80 or < −0.80), with polynomial regression trend lines for positively and negatively correlated groups. **f**, Normalized gene expression (Z-score) of select highly correlated genes. **g**, Expression of *AIL6* and *RIL1* in W2.5 spikes. **h**, Expression domains annotate and group leaf ridges (LRs) from 1 (basal) to 13 (apical). **i**, PC analysis of averaged transcripts per LR group. **j**, Normalized gene expression (Z-score) of highly correlated genes to PC1 (> 0.80 and < −0.80), with polynomial regression trend lines shown separately for positively and negatively correlated groups. **k**, Normalized gene expression (Z-score) of select highly correlated genes. **l**, Expression of *YABBY1* and *SPL17* expressed in W2.5 spike.

To quantify these gradients, we grouped cells from each SR and LR, ordered them longitudinally from basal (1) to apical (13; Fig. 5c,h), and performed principal component (PC) analysis using average normalised counts per cell in each group (Fig. 5d,I; Supplementary Table 15). PC analyses revealed a consistent developmental progression for SRs and LRs along PC1 (Fig. 5d). Genes positively correlated with PC1, such as *KNOX5* (0.867), *AINTEGUMENTA-LIKE6* (*AIL6,* 0.877), and *INCREASED LEAF INCLINATION 3 (ILI3,* 0.828), have elevated expression in more central/apical SRs (Fig. 5f,g; Supplementary Table 16). Negatively correlated genes, including *SQUAMOSA-PROMOTER BINDING PROTEIN-LIKE 14* (*SPL14,* - 0.890), *FRUITFULL3* (-0.820), and *RACHIS-LIKE1* (*RIL1,* -0.923), were more prominent in basal SRs (Fig. 5f; Supplementary Table 16). Leaf ridges showed a similar separation along PC1; *APETALA2-5* (-0.932), *SPL17* (-0.889), and *LEAFY COTYLEDON 1 (LEC1,* -0.934) showed elevated expression in basal LRs, contrasting to *YABBY1* (0.944) and *MADS32* (0.894) with higher expression in more central/apical LRs (Fig. 5k,l; Supplementary Table 16). Thus, based on the highly correlated genes across SRs and LRs, we observed two opposing gradients along the apical-basal axis, which comprise different sets of genes, but which cross at a mirrored position along the inflorescence (SR5 & LR6; Fig. 5e,j). This suggests that adjacent spikelet and leaf ridges coordinate gene expression gradients, and collectively shift their specific expression profiles at the position where SM first start to actively differentiate.

## Discussion

A systematic and unbiased approach is critical to uncover biologically meaningful insights from spatial transcriptomics data. To achieve this, we clustered cells across all developmental stages into EDs and identified enriched gene markers. By concatenating samples, we traced EDs across time, revealing novel insights into tissue identity and differentiation (Fig. 2). For example, a small population of three ED15 cells identified at W3.25 marked the earliest palea progenitors, a discovery that would have been missed through single-sample clustering. EDs also revealed similarities between tissue types. Both developing leaves at W2.5 and lemmas at W5 consist of ED1 and ED2, supporting the classification of lemmas as leaf-like organs, rather than floral-tissues^31^. Concatenating samples before clustering provided novel biological insights, therefore, we propose this strategy to analyse spatial transcriptomics data across developmental time-courses.

By utilising expression domains, we characterised each phytomer unit along the apical-basal axis, capturing the transcriptional programmes patterning vegetative and inflorescence tissues. At W2.5, ED9 cells marked by *MND1*, were identified in axillary meristems below the inflorescence (Fig. 3). These cells, forming in the axils of leaves (ED1,2,13,9), are specific to the transcriptional program driving leaf outgrowth and axillary meristem suppression. In contrast, the inflorescence is patterned by LRs (ED12) and SRs (Fig. 5), with SR adaxial boundaries consistently marked by *CUC3* and *BARREN STALK1* (ED11)^32–35^. However, SR patterning also varied along the apical-basal axis. Basal SRs (SR1–5) show increased ED12-type cells and higher expression of genes involved in AM suppression like *MND1*^36^, while central and apical SRs (SR6–13) expressed meristem-specific genes like *KNOX5*^29^ and floral identity-promoting factors such as *AIL6*^37,38^. Additionally, the cellular resolution of MERFISH revealed not only differences between meristems but also within them, uncovering spatially restricted A*IL6* and *ILI3* expression along the adaxial-abaxial axis (Extended Data Fig. 6).

Gene expression patterns distinguishing SRs form opposing gradients that define their developmental progression, as revealed by PC analysis (Fig. 5). A similar apical-basal gradient emerged in LRs. While all LRs clustered into ED12, basal LRs specifically expressed additional bract suppression genes (*SPL17/TSH4* and *RIL1*)^8,39^, suggesting a more complex LR suppression network at the base of the spike involving factors beyond *TSH1*. Models of inflorescence development propose that the suppressed LR serve as a signalling centre orchestrating neighbouring SR meristem activity^40^. The fined-tuned transcriptional patterns of basal LRs and SRs, which are unique compared to central/apical regions, support their coordinated activity. Importantly, this insight into their spatial coordination is only achievable with spatially resolved techniques, which overcome the limitations of bulk tissue or single-nuclei RNA sequencing.

We hypothesize that the opposing gradients observed along the inflorescence may define meristem phase transition^41^. Prusinkiewicz et al. proposed that meristem transitions are governed by the degree of ‘vegetativeness’, which must drop below a threshold to enable flowering^42^. Backhaus et al. put forward a wheat-specific model to meristem transition in which basal phytomers, initiated shortly after the SAM transitions to an inflorescence meristem, encounter heightened ‘vegetative’ signals, which may account for reduced SR development and enhanced LR growth^13^. We suggest that the distinct yet spatially coordinated transcriptional states of basal SRs and LRs adds to this understanding, potentially highlighting novel factors involved in meristem transition.

In summary, applying the cellular resolution of MERFISH to plant tissues allowed us to map the expression of a curated 200-gene panel. While the panel is limited in the breadth of genes assayed, our systematic approach combining sample integration, expression domain clustering, and gene enrichment analysis, provided novel candidate factors contributing to apical-basal patterning in wheat spikes. To support further research, all raw data is made available in addition to analysed data accessible via a WebAtlas^21^ interface (www.wheat-spatial.com). We aim to empower the research community to leverage spatial transcriptomics analyses as a valuable tool in plant biology.

## Author Contributions

**K.A.L.**: Conceptualization, Data Curation, Formal Analysis, Investigation, Methodology, Project Administration, Resources, Software, Validation, Visualization, Writing – Original Draft Preparation; **A.L.**: Investigation, Data Curation, Methodology, Platform management, Project management, Data management, Validation, Visualization; **M.R.W.J.**: Data Curation, Formal Analysis, Investigation, Resources, Writing – Original Draft Preparation; **N.M.A.**: Resources, Writing – Original Draft Preparation, Writing – Review & Editing; **R.E.E.**: Data Curation, Resources, Software, Visualization; **C.C.**: Methodology, Resources; **S.J.C.**: Formal Analysis, Validation, Writing – Review & Editing; **X.M.L.**: Resources; **A.E.B.**: Resources, Writing – Review & Editing; **A.G.**: Resources, Writing – Review & Editing; **V.K.**: Methodology, Resources; **Y.P.**: Resources; **M.V.**: Resources, Software, Supervision; **B.S.**: Resources, Software, Supervision; **G.G.K.**: Data Curation, Software; **J.X.**: Resources; **W.H.**: Conceptualization, Funding Acquisition; **I.C.M.**: Funding Acquisition, Methodology, Project Administration, Supervision; **C.U.**: Conceptualization, Formal Analysis, Funding Acquisition, Methodology, Project Administration, Supervision, Visualization, Writing – Original Draft Preparation.

## Acknowledgments

The authors would like to thank Susan Duncan (JIC) for help with training in sample preparation, David Wright (EI) and David Swarbreck (EI) for supporting data management discussions, and Philippa Borrill (JIC) and Chen Ji (JIC) for contributing grain-related genes for panel design. This work was supported the European Research Council (ERC-2019-COG-866328) and the UK Biotechnology and Biological Sciences Research Council (BBSRC) through the Delivering Sustainable Wheat (BB/X011003/1), Building Robustness in Crops (BB/X01102X/1), and Cellular Genomics (BB/X011070/1) Institute Strategic Programmes. This work was delivered via Transformative Genomics, the BBSRC funded National Bioscience Research Infrastructure (BBS/E/23NB0006) at EI by members of the Single-Cell and Spatial Analysis Group and Technical Genomics Group. K.A. L. was supported by the Gatsby Charitable Foundation; S.J.C was supported by the UKRI Biotechnology and Biological Sciences Research Council Norwich Research Park Biosciences Doctoral Training Partnership (BB/T008717/1); and J.X was supported by the Beijing Natural Science Foundation Outstanding Youth Project (JQ23026).

## Code and Data Availability

Code associated with this project is available at Github including all implementation scripts (https://github.com/katielong3768/Wheat-Inflorescence-Spatial-Transcriptomics). All files, images, and segmentation outputs used in the analyses and described in GitHub are available on an individual sample basis (https://zenodo.org/records/14515927). The RNA-sequencing data generated in this study were submitted to NCBI under BioProject number XXXXX. Visualisation of transcript counts per cell for *P1*^WT^ samples are available at www.wheat-spatial.com.

## Methods

### Plant materials

We used two BC_6_ near isogenic lines (NIL) differing for *VRT-A2* alleles in a hexaploid wheat (cv Paragon) background. One NIL carried the wildtype Paragon *VRT-A2a* allele, here named *P1^WT^*, whereas the second NIL carried the *VRT-A2b* allele from *Triticum turgidum* ssp. *polonicum* (named *P1^POL^*)^14^. Plants were grown under a 16/8 h light/dark cycle at 20/15 °C, 65% relative humidity and bottom-watering irrigation^43^.

### Dissections and Sample Preparation

The *VRT-A2a* NIL was used for semi-spatial RNA-seq, whereas both NILs were used for MERFISH. For semi-spatial RNA-seq we used a published dissection methodology^44^ to produce basal and central/apical sections. At the Early Double Ridge stage (EDR, W2), spikes were bisected, whereas for the Late Double Ridge (LDR, W2.5), Lemma Primordia (LP, W3.25), Terminal Spikelet (TS, W4) and Carpel Extension (CE, W5) stages, the basal section consisted of the most basal four spikelets from each spike. Two spikelets were skipped, then the subsequent four spikelets were harvested to comprise the central section (Extended Data Fig. 1; Supplementary Table 1). Samples were stored at -70 °C until RNA extractions which were conducted from the pooled microdissected spikes using Qiagen RNeasy Plant Mini and Zymo Direct-zol RNA Microprep kits as described in the manufacturer’s manual. Total RNA (1 ug) was sent to Novogene UK for PCR-free library preparation and Illumina sequencing (PE150; 50M reads per sample).

For MERFISH, we used a similar dissection protocol^44^, but maintained the youngest leaves surrounding meristems (Supplementary Fig. 1). After dissection, meristems were transferred using an RNase-free pipette tip into 4% paraformaldehyde (PFA) in 1× PBS (prepared from 6% formaldehyde [w/v], methanol-free; Pierce 28906) in 2 mL RNase-free Eppendorf tubes. Samples were vacuum infiltrated for 10 minutes or until tissue sank and incubated overnight at 4 °C. The PFA solution was removed, and the samples were washed three times with 1× PBS. Tissue was then immersed in 15% sucrose in 1× PBS at 4 °C for 6 hours, followed by immersion in 30% sucrose in 1× PBS at 4 °C overnight.

### Analysis of semi-spatial RNA-seq data

We trimmed raw reads with cutadapt (v1.9.1)^45^ and generated read counts and transcripts per million (TPM) values using Kallisto pseudo-alignment (v0.44.0)^46^ for all genes in the IWGSC RefSeq v1.1 annotation^47^ (Supplementary Table 1). We conducted subsequent analyses for high confidence gene models with non-zero counts in at least one sample. We transformed read counts (rlog function; DESeq2 (v1.34.0))^48^, and performed principal component (PC) analysis with prcomp^49^; we identified no outliers (Extended Data Fig. 1b). We calculated differential expression (p < 0.001; Benjamini-Hochberg corrected) between central and basal sections across time using ImpulseDE2 (v 3.6.1)^50^, on genes with average > 0.5 TPM for at least one stage-section combination (Supplementary Table 2). We clustered the 12,384 differentially expressed genes using k-means (k1:10) and displayed with pheatmap (v1.0.12)^51^.

### Gene Panel Selection and Design for MERFISH

We designed a 300-gene panel for MERFISH, comprising 200 genes associated with spike development (116 from differential expression dataset, 73 additional genes, and 11 housekeeping genes), and 100 genes from a separate wheat grain project which are not described here. We removed genes that could not accommodate at least 25 specific probes, based on Vizgen’s probe design software, except for three genes targeted by >20 probes. MERFISH probes were designed and synthesized by Vizgen (Supplementary Table 3).

### Meristem Embedding and Sectioning

We cleaned all surfaces and dissection tools with RNABlitz before use. We marked a 1 cm × 1 cm area on the back of a Tissue-Tek mold (25 × 20 × 5 mm; Thermo Fisher, AGG4580) and filled with Tissue-Plus OCT compound (Agar Scientific, AGR1180). We also filled a 60 mm Petri dish with OCT. We removed individual meristems from the 30% sucrose solution using clean dissection tools, aided by a drop of OCT on the tool to adhere the meristems during collection. We transferred meristems to the OCT-filled Petri dish, where they were mixed with OCT to remove residual sucrose and ensure complete coating. Using a stereomicroscope (Leica S9 with an HXCAM HiChrome HR4 Lite camera and a Photonic Optics light source), we inspected meristems for air bubbles, which were carefully removed with a fine dissection tool. We trimmed excess vegetative tissue as needed. Meristems were then placed into the OCT-filled Tissue-Tek mold, arranged within the marked 1 cm^2^ region according to genotype and developmental stage. Each OCT block contained 5–36 meristems, depending on the developmental stage (Supplementary Fig. 1), and we imaged them using GX Capture-T^52^. The OCT blocks were flash-frozen and stored at −70 °C.

We performed sectioning using a Leica CryoStar NX70. All inside surfaces and tools were cleaned with Blitz RNase Spray (Severn Biotech Ltd, 40-1735-05), and a fresh blade (MX35 Ultra™ Microtome Blade, 3053835) was used. We set the chuck temperature to −20 °C, and the blade temperature to −18 °C. We pre-warmed samples at the back of the cryostat for 30 minutes before sectioning, and brought the MERSCOPE Slides (Vizgen, 20400001) to room temperature. We trimmed OCT blocks to remove excess OCT, mounted to the chuck, and further trimmed until tissue was exposed; 10 µm sections were cut to inspect tissue regions on glass slides. Once we identified the region of interest at the optimal depth and angle, 10 µm sections were flattened with paintbrushes, flipped, and mounted onto room-temperature MERSCOPE slides, following the placement and technique outlined in the MERSCOPE user guide. After mounting, we placed the slides in 60 mm Petri dishes and incubated at the back of the cryostat for 30 minutes. We then fixed the slides in 4% methanol-free PFA in 1× PBS for 10 minutes. We washed the slides three times with 1× PBS, incubating for 5 minutes per wash. We aspirated residual PBS and air-dried the slides for 1 hour in a cell culture hood with the Petri dish lid closed. We then incubated the slides with 5 mL of 70% ethanol prepared in RNase-free water. Petri dishes were sealed with parafilm and stored at 4 °C, either overnight or for up to 7 days.

### MERSCOPE Workflow

We performed slide preparation following guidelines for non-resistant fixed frozen tissue clearing (91600002_MERSCOPE Fresh and Fixed Frozen Tissue Sample Preparation User Guide_Rev E (Vizgen)). We prepared slides using Vizgen sample preparation kit (Vizgen, 10400012) with all instruments (including hybridisation box (Brabantia, 203480)) cleaned using both 70% ethanol and then RNAseZAP (Invitrogen, AM9782) or Blitz RNase Spray (Severn Biotech Ltd, 40-1735-05). We performed checks for autofluorescence at 10X using an EVOS FL2 microscope under a DAPI light cube, recording light intensity levels to decide on reduction of autofluorescence before and after photobleaching (performed for between 3 and 8 hours in EtOH 70%, Vizgen 10100003). A 300-gene probe set (Vizgen product number 20300007) was applied and hybridised for 48 hours. On the days of a run, we re-checked autofluorescence levels and topped up using the photobleacher for 3 hours if necessary. After DAPI staining, we also made checks for efficiency of staining. Clearing times varied depending on run slots. A standard clearing at 47 °C for 1 day with clearing buffer (Vizgen) containing proteinase K (NEB, P81070S) addition, was always performed, whereas additional days (1 to 4 days) of clearing at 37 °C without proteinase K in the buffer was performed. Tissue never fully cleared by eye nor when using a light microscope before MERSCOPE runs.

Upon imaging, care was taken to minimise smears and lint on the slide by cleaning with 80% ethanol and lens cleaning tissue (2105-841, Whatman). We outlined regions of interest around individual spikes on the slide overview using DAPI staining (Supplementary Figure 1). Following 60x imaging, we decoded transcripts using the panel specific MERSCOPE Codebook. We processed raw data with the MERSCOPE Instrument Software to generate and output file structures as described in MERSCOPE instrument User Guide.

### Cell segmentation and processing

We performed cell segmentation on stitched images of DAPI and PolyT staining. Prior to segmentation and to minimize error, we lightened seam lines in the stitched images using FIJI (version 1.54f)^53^. Dark stitching lines were processed using the following steps: ‘Process>Filters>Maximum’ (radius = 2 pixels, applied twice) and ‘Process>Filters>Median’ (radius = 2 pixels, applied twice). Then applied three times across the entire image: ‘Process>Filters>Gaussian Blur’ (radius = 4 pixels). See Supplementary Fig. 4 for image edits and segmentation results.

We performed cell segmentation and transcript assignment using the Vizgen Post-Processing Tools (VPT, version 1.2.2)^23^, within a Python virtual environment on Ubuntu 20.04. We used Cellpose2 cyto2 model^17^ with DAPI (blue channel) as the nuclear marker and PolyT (green channel) as the cytoplasmic marker. For segmentation parameters, see logs (https://zenodo.org/records/14515927). We exported segmentation results as polygon geometry in both mosaic and micron space, assigned transcripts to cell boundaries using the partition-transcripts function in VPT, and generated cell metadata with the derive-entity-metadata function. Finally, we integrated cell boundaries into existing .vzg files for visualization in the Vizgen MERSCOPE Visualizer Tool^54^ using the update-vzg function. All implementation scripts are available (https://github.com/katielong3768/Wheat-Inflorescence-Spatial-Transcriptomics) with example commands (https://zenodo.org/records/14515927).

### Quality Checks and Filtering

We loaded the spatial transcriptomic data from the eight samples into AnnData objects (anndata v0.10.7)^55^ and processed using Squidpy (v1.4.1)^19^ and Scanpy (v 1.10.0^18^. We filtered expression data to include the 200 spike development and control genes, selected cells from a single inflorescence within the imaged area, and excluded low-quality cells based on volume (>500 pixels) and transcript count (>25 counts)(Supplementary Table 4). For each sample we calculated quality control (QC) metrics (total counts per cell, number of genes detected per cell, percentage of counts from top-expressed genes) and summary statistics (total counts per cell, detected genes per cell; Supplementary Table 5; Supplementary Fig. 2). We assessed off-target hybridization using blank probes (Supplementary Table 6), and across wheat homoeologs (Supplementary Note 1; Extended Data Fig. 2). Finally, we normalised expression data for all samples using scanpy functions sc.pp.normalize_total() and sc.pp.log1p() (v 1.10.0)^18^.

As a further quality control metric, we performed *in silico* spike dissections equivalent to those captured by physical microdissection for 16 high-quality MERFISH sections using the MERSCOPE Visualiser ‘draw ROI polygon’ tool^54^. We exported frequency tables of MERFISH transcripts within basal and central/apical regions and calculated Spearman’s rank correlation coefficients between these values and the mean TPM values from the relevant genotype-section-stage combination of the semi-spatial RNA-seq data (Supplementary Table 7; Supplementary Fig. 2).

Additionally, we identified *in situ* hybridisation results in wheat, barley, rice, and maize from equivalent tissues and time points as those used for MERFISH (Supplementary Table 8) and visualised them in side-by-side comparisons (Supplementary Fig. 3). We also compared expression profiles of genes involved in the ABCDE model of floral development with mutant phenotypes and knowledge from orthologs in rice, maize, barley, and wheat (Supplementary Note 2; Supplementary Table 12), and visualised using www.wheat-spatial.com (Extended Data Fig. 5).

### Cell Segmentation and Transcript Visualisation

We processed the cell segmentation data as GeoDataFrames using the GeoPandas (v0.14.4)^56^, and converted the transcript coordinates into a GeoDataFrame from global x and y coordinates. We performed a spatial join operation to assign transcripts to segmented cells, retaining only transcripts located within cell boundaries. We next rotated segmented cell polygons and transcript coordinates using NumPy (v1.26.3)^57^, and visualised cell geometries as polygons using Matplotlib (v 2.0.4)^58^ with polygon handling and transformations facilitated by Shapely (v2.0.4)^59^. Transcripts were overlaid as point features (Fig. 2b-e,g-h, 3a-e, 4a-b, 5a-c,g-h,l). Full details in implementation scripts (https://github.com/katielong3768/Wheat-Inflorescence-Spatial-Transcriptomics).

### MERFISH Data Integration, Unsupervised Clustering, and Gene Enrichment Analysis

We processed spatial transcriptomic data from eight samples (four timepoints; two NILs) using the Scanorama (v 1.7.4)^60,61^ integration tool, and performed clustering using the Leiden algorithm with a resolution parameter of 1.0 (Fig. 2a). Spatial maps of Leiden cluster assignment were performed as described in ‘Cell Segmentation and Transcript Visualisation’ (Fig. 2b-e; Extended Data Fig. 3). We exported the expression domain (ED) assignment of each per cell (Supplementary Table 9), and visualised as percentage of cells in each ED, per sample (Fig. 2f). Next, we performed gene enrichment analysis on the integrated AnnData object with Scanpy function sc.tl.rank_genes_groups() using the logistic regression model^20^. This analysis returned a ranked list of genes most probable to be enriched gene markers, which we displayed alongside the average normalised expressions per ED for each sample (Supplementary Table 10). We determined top enriched values (using a +2 standard deviation threshold) and used these to annotate EDs with tissue type identity labels (Supplementary Table 11).

### Transect analysis of *VRT-SEP* gradients

We filtered cells from two samples (W4, *VRT-A2a* and *VRT-A2b NILs*) to include only cells from the inflorescence region, defined as the beginning of ED12 marking leaf ridges. These cells were selected in the MERSCOPE Visualizer tool^54^ using the Polygon Lasso Tool, exported as a.csv file, and the segmented cells and transcripts were mapped as previously described. The Y-axis of the spatial plot was divided into 25 transverse bins along the spike. Each cell was assigned to a bin based on its centre Y-coordinate, and we averaged the normalized transcript counts per cell within each bin (Fig. 4a-b). For both samples, we binarized gene expression data for *VRT2* and *SEP1-4* within each cell, assigning a value of 1 for detected reads and 0 for no detected reads. For each ED, we quantified the number of cells expressing only *VRT2*, only *SEP1-4*, or co-expressing both genes and visualised them as a percentage with Matplotlib (v 3.8.2^58^; Supplementary Table 13; Fig. 4c,d).

### Gene expression analysis on Late Double Ridge Spikes

We selected Late Double Ridge (W2.5) and Lemma Primordia (W3.25) *P1^WT^* inflorescence cells using the MERSCOPE Visualizer Polygon Lasso tool and exported cell identity data as a .csv file. We defined the inflorescence boundary by the first suppressed leaf ridge (ED12) and excluded cells outside the inflorescence (Supplementary Table 14). Cell counts were summed by ED, and we calculated the cumulative percentage of cells in the most populated ED to assess their contribution to the total cell population. We calculated the top Eds accounting for approximately 94% of the cells in the sample.

Groups of cells comprising the Leaf Ridges (LR) and Spikelet Ridges (SR) (defined by EDs) were annotated as “Custom Cell Groups” in the MERSCOPE Visualizer tool^54^. We delineated SR boundaries by ED4 cells along the adaxial and abaxial axes, extending to the start of ED3 cells along the medio-lateral axis. We identified LRs as groups of ED12 cells beginning beneath the end of ED4 cells from adjacent SRs. We labelled SRs and LRs sequentially from 1 (most basal) to 13 (most apical) along the inflorescence (Fig. 5c,h; Supplementary Table 14). We calculated the total number of cells in basal (LR1-4) and central (LR8-11) leaf ridges and determined the mean cell numbers and summary statistics to compare ridge sizes between these regions.

Normalized gene expression values per cell were averaged by LR or SR group and filtered to include only genes with at least one average expression score above 0.30 across all groups (Supplementary Table 15). We standardised the resulting data matrix using StandardScaler and performed PC analysis with scikit-learn (v 1.4.2)^62^, to extract the first two principal components (Fig. 5d,i). We inverted PC1 and PC2 scores to align the axes with the desired biological orientation, and calculated correlations between individual genes and PC1 (Supplementary Table 16).

We selected genes with strong positive or negative correlations to PC1 (>│0.80│). We calculated average expression (Z-score normalised) trends for these groups and smoothed values using a Savitzky-Golay filter (SciPy v1.13.0)^63^. Individual gene trends were also smoothed for visualisation. The smoothed average trends were fitted with cubic polynomial curves (NumPy v 1.26.3)^57^ to highlight overall expression gradients (Fig. 5e,j). Plots were generated using Matplotlib (v3.8.2)^58^ and Seaborn (v0.13.1)^64^, and all scripts are available (https://github.com/katielong3768/Wheat-Inflorescence-Spatial-Transcriptomics).

Next, we developed maps of gene expression inside spikelet meristems focused on SR8–11. We selected individual spikelet meristems using predefined cell groups and rotated the cells to achieve a consistent orientation across the four SRs. We filtered transcript coordinates to include only those falling within selected cells and defined a 25 × 25 grid over the SR cells. We calculated transcript counts for each gene within each grid bin using NumPy digitize function (v1.26.3)^57^, resulting in frequency matrices for genes of interest . We then averaged frequencies across SR8-11 to generate composite frequency matrices for each gene. To enhance spatial patterns, we applied smoothing to the composite matrices using a Gaussian filter with SciPy (v1.13.0)^63^. To suppress low-frequency noise, we also applied thresholding by setting the bottom 20% of values as minimum the threshold. Finally, we generated contour plots to visualize smoothed composite matrices (Matplotlib, v3.8.2)^58^.

