## Extended Data Figure for "Spatial Transcriptomics Reveals Expression Gradients in Developing Wheat Inflorescences at Cellular Resolution"

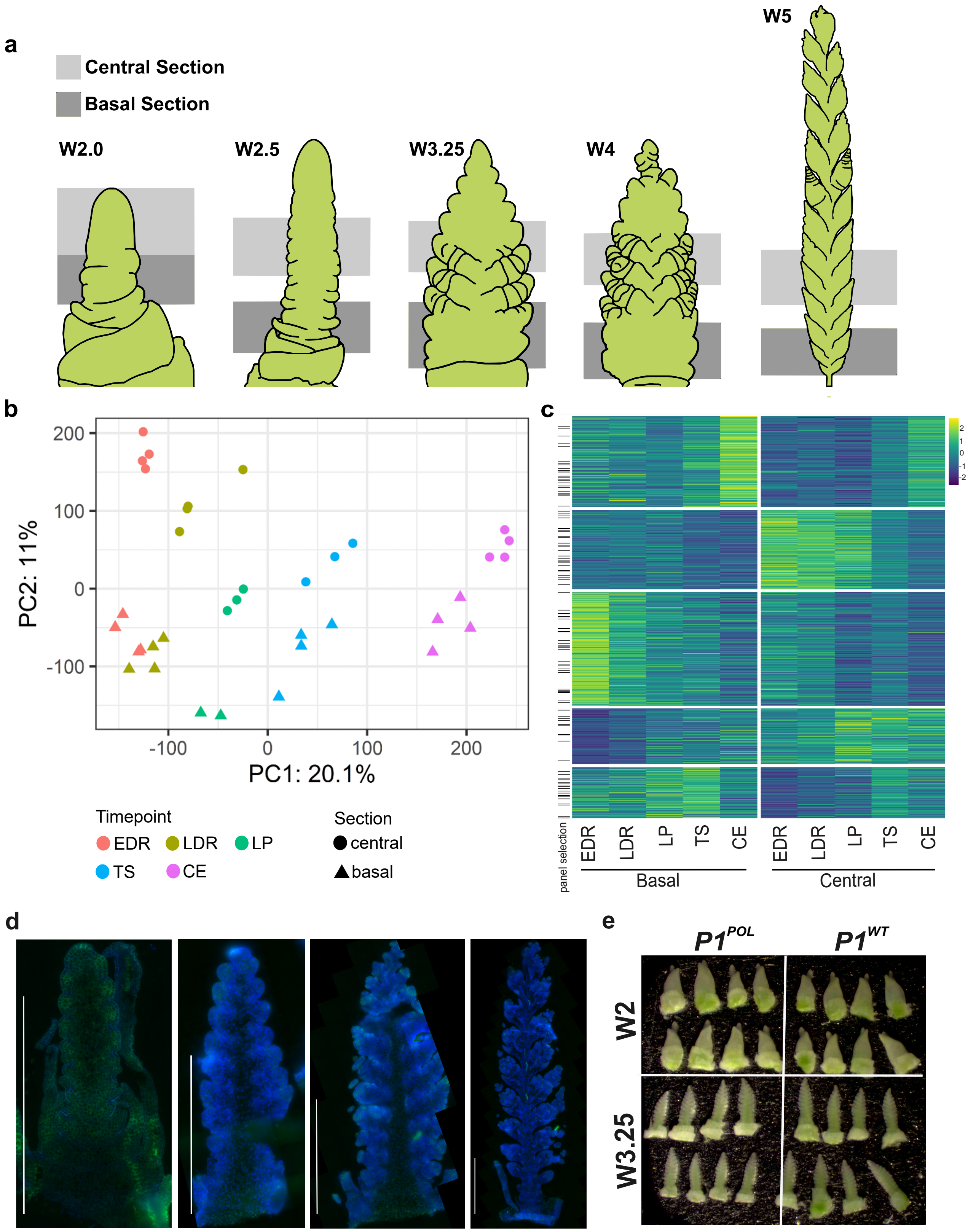


**Extended Data Figure 1. Micro-dissection of wheat inflorescence and pooled RNA-sequencing distinguishes samples by section and stage.**

**a,** Location of central and basal sections for each developmental time point used in pooled tissue RNA-sequencing. **b,** Principal component (PC) analysis separate samples by timepoint (PC1) and spatial section (PC2). EDR, Early Double Ridge; LDR, Late Double Ridge; LP, Lemma Primordium TS, Terminal Spikelet; CE, Carpel Extension. **c,** 12,384 genes differentially expressed between central and basal microdissections of wheat inflorescences across five developmental time points, genes selected for panel annotated with black bars. **d,** Representative cryosections of four developmental stages: Late Double Ridge (LDR, W2.5), Lemma Primordia (LP, W3.25), Terminal Spikelet (TS, W4), and Carpel Extension (CE, W5); DAPI staining in blue, PolyT in green, visualised with MERSCOPE Visualizer Tool. **e,** OCT-embedded block of 32 dissected meristems from stages EDR and LP, representing *P1*$^{WT}$ and *P1*$^{POL}$ genotypes.

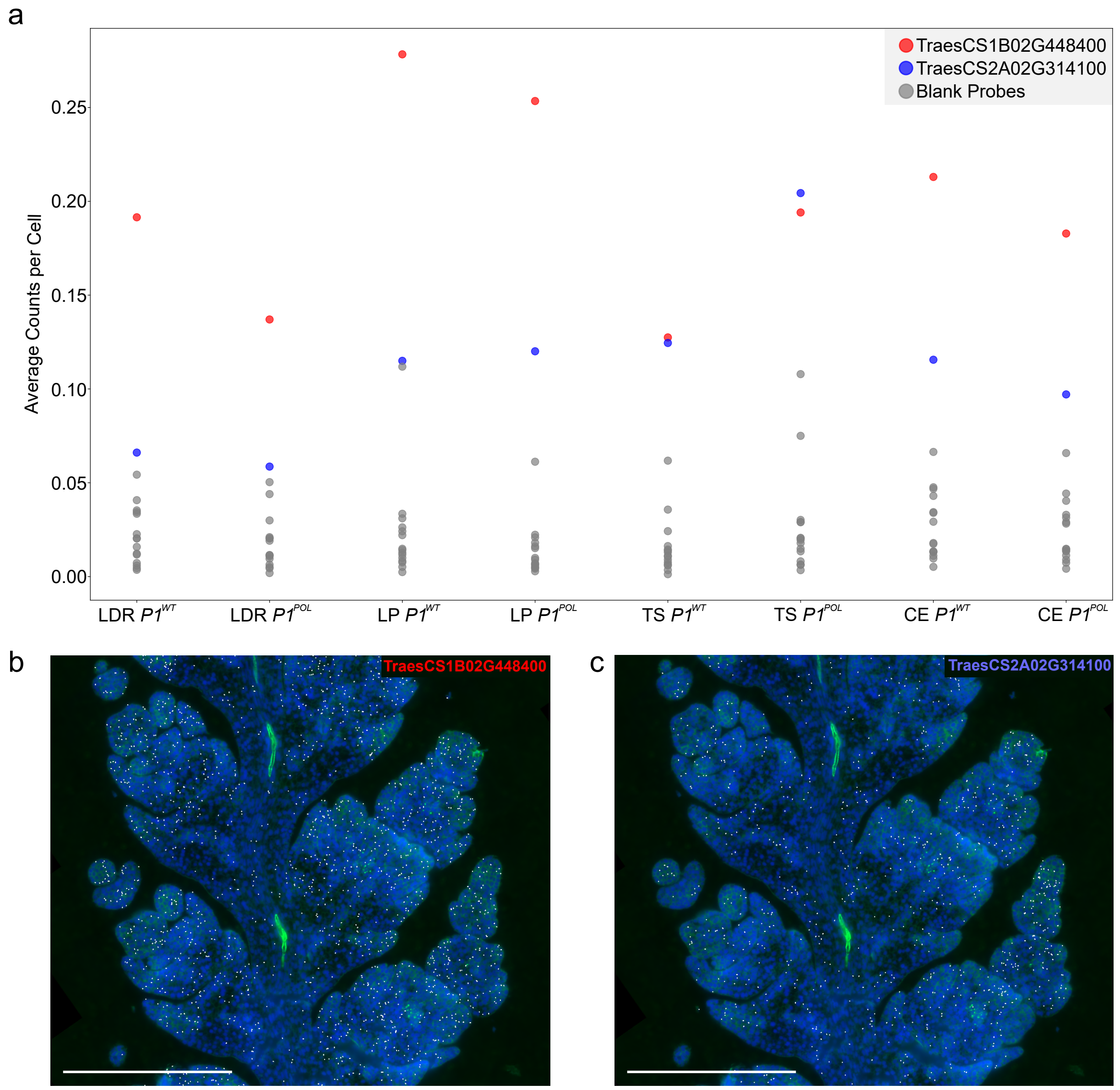


**Extended Data Figure 2. Hybridisation patterns indicate non-homoeolog specificity and low off-target rates for MERFISH probes.**

**a,** Average count per cell metrics of 15 blank probes (grey) and two gene encoding probes, *TraesCS1B02G448400* and *TraesCS2A02G314100*, designed to test off-target activity, as described in Supplementary Note 1. **b,c,** Expression patterns in W5 spikes of gene encoding probes (**b**) *TraesCS1B02G448400* and (**c**) *TraesCS2A02G314100* visualised in the MERSCOPE Visualizer Tool. DAPI stain in blue and detected transcripts in white.

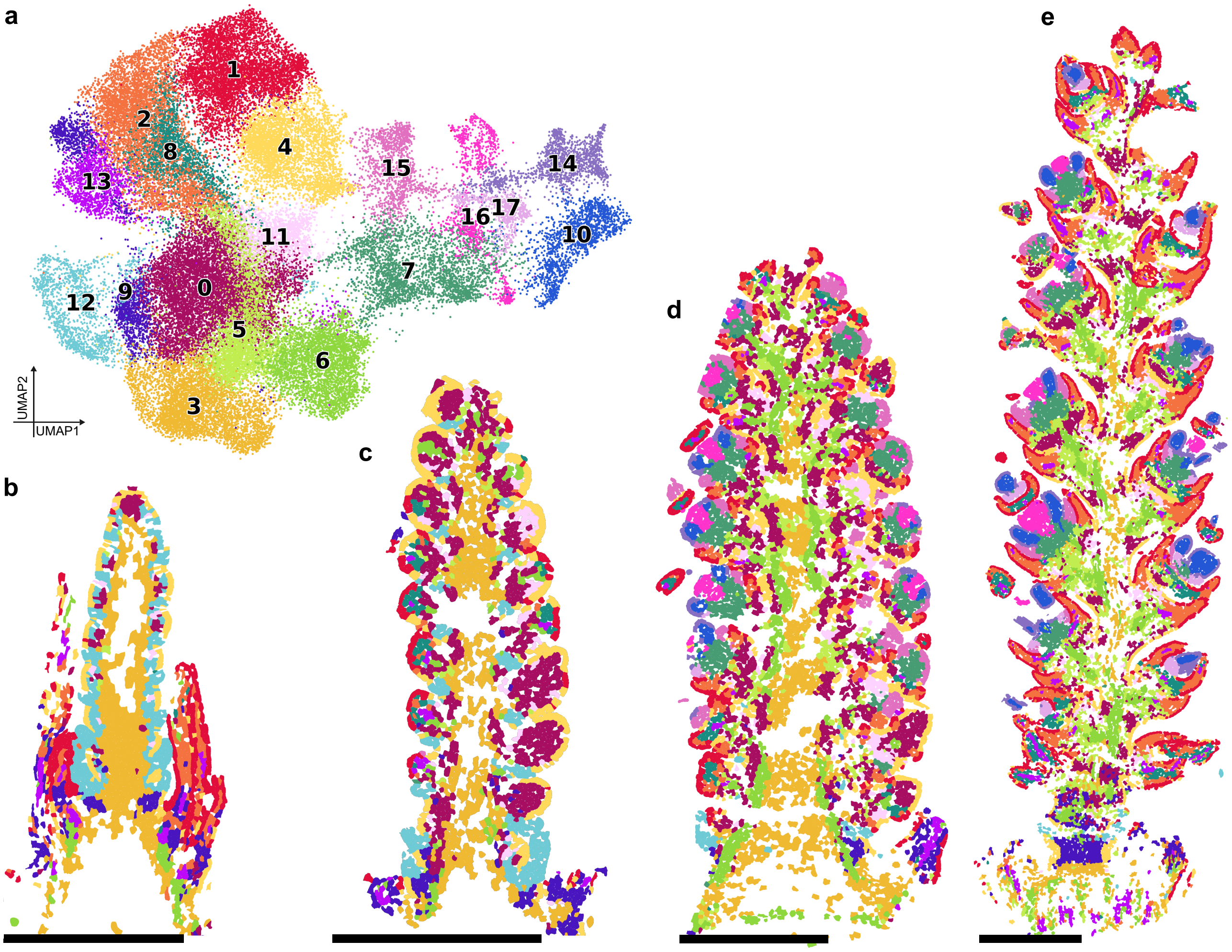


**Extended Data Figure 3. Spatial maps of 18 expression domains mapped over four developmental stages in *P1^POL* NIL replicates.**

**a,** UMAP projection of cells from eight samples, and expression domain assignment as in Fig. 2a. **b-e,** Spatial maps of Leiden clustering across time points (**b**) W2.5, (**c**) W3.25, (**d**) W4, and (**e**) W5 using Squidpy (v1.4.1), Scanpy(v 1.10.0), and Scanorama(v1.7.4). Scale bar = 500 µm.

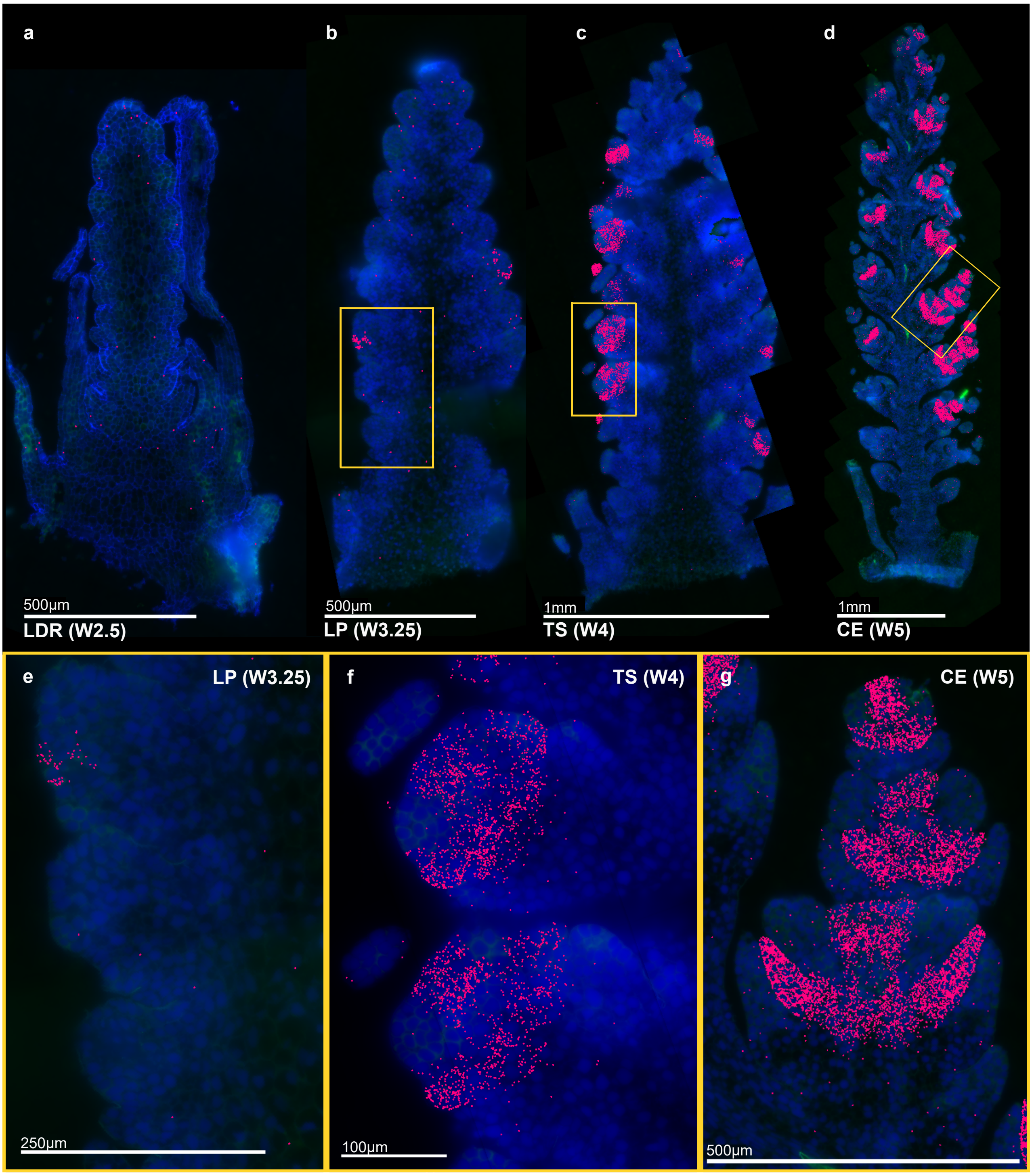


**Extended Data Figure 4. Spatial expression patterns of *AGL6* marks palea and palea progenitor cells.**

**a,d,** Expression patterns of *AGL6* in wildtype spikes at (**a**) LDR/W2.5, (**b**) LP/W3.25, (**c**) TS/W4, and (**d**) CE/W5 developmental stages. DAPI channel in blue, *AGL6* transcripts in pink, visualised with the MERSCOPE Visualizer Tool. LDR, Late Double Ridge; LP, Lemma Primordium TS, Terminal Spikelet; CE, Carpel Extension. **e-g,** Higher magnification of *AGL6* expression at (**e**) LP, (**f**) TS, (**g**) CE stages.

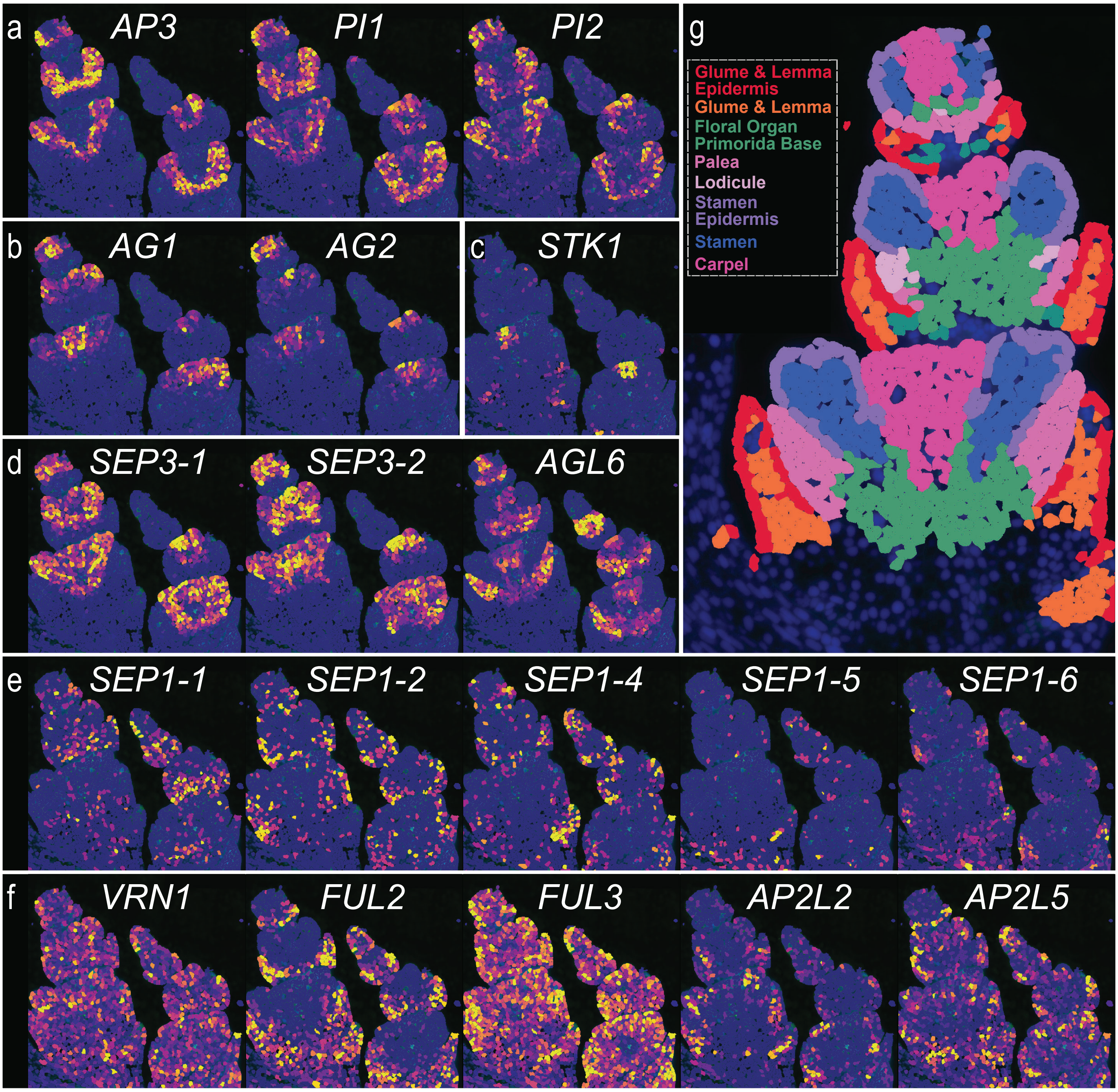


**Extended Data Figure 5. Spatial expression patterns of putative floral organ identity genes.**

To test the assumptions of the ABCDE model in wheat, we use MERFISH to investigate spatial expression patterns of (**a**) B-class genes *AP3*, *PI1*, and *PI2*, (**b**) C-class genes *AG1* and *AG2*, (**c**) D-class gene *STK1*, and (**d**) E-class genes *SEP3-1*, *SEP3-2*, and *AGL6* in W5 florets. The genes in **a-d** are expressed in a highly spatially defined manner, suggesting their role as floral organ identity genes as postulated by the ABCDE model. In contrast, genes belonging to (**e**) the *LOFSEP* clade or (**f**) encoding putative A-class function (*AP1/FUL-like* or *AP2-like*) show either little or unrestricted expression in developing florets, suggesting that they do not contribute to defining floral organ identity. **g,** Expression domains (EDs) obtained through cell segmentation and unsupervised clustering accurately predict floral organ identity. This image corresponds to the left-hand spikelet of each panel in **a-f.** Note that the heatmaps in **a-f** are not scaled uniformly, but instead to maximize visibility of each gene's expression domain. Hence, transcript levels in these images are not comparable. For further details see www.wheat-spatial.com.

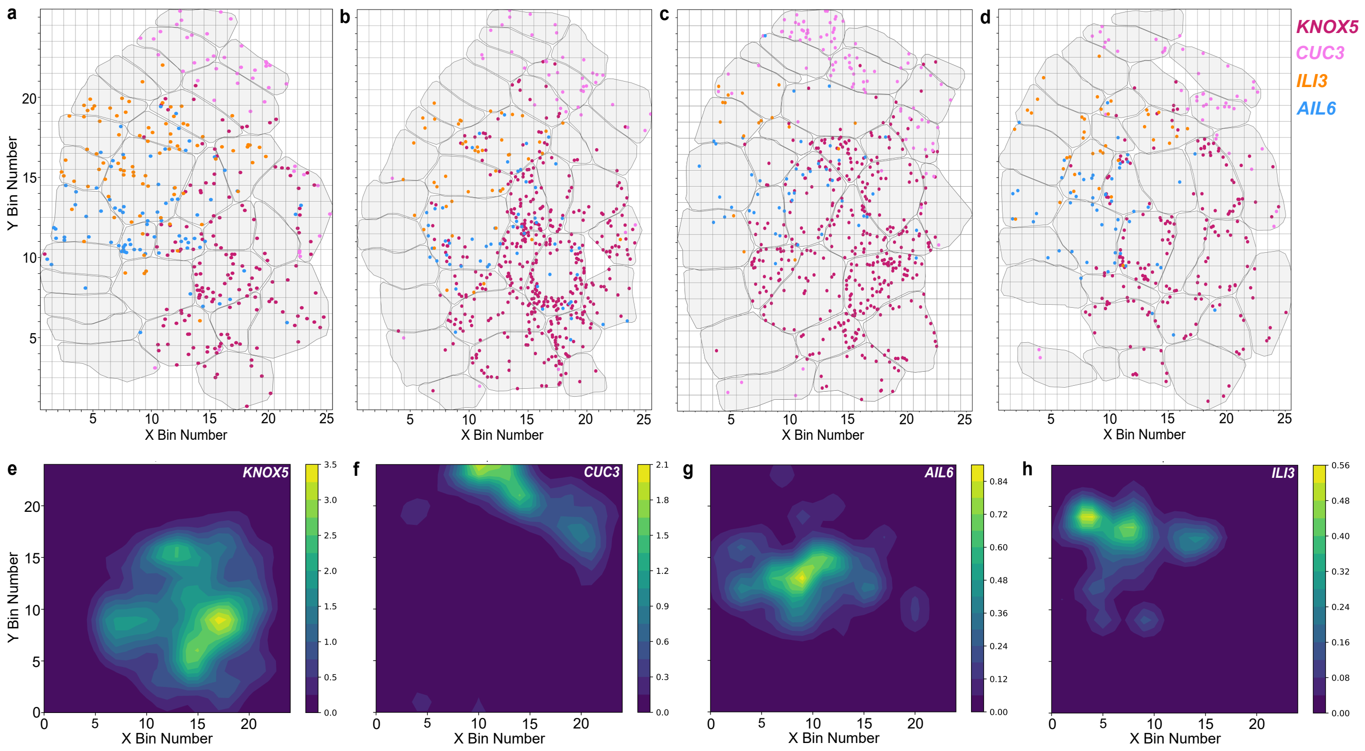


**Extended Data Figure 6. Composite maps of gene expression within spikelet ridges reveals phased gene expression patterns across meristems.**

**a,** Individual SR8 meristem spatial map with overlaid expression of *KNOX5*, *CUC3*, *ILI3*, and *AIL6*. **b-d,** Equivalent spatial maps for SR9-11, respectively. **e-h,** Composite maps of average expression from SR8-11 summarised for genes (**e**) *KNOX5*, (**f**) *CUC3*, (**g**) *AIL6*, and (**h**) *ILI3*.
