## Supplementary Figures for "Spatial Transcriptomics Reveals Expression Gradients in Developing Wheat Inflorescences at Cellular Resolution"

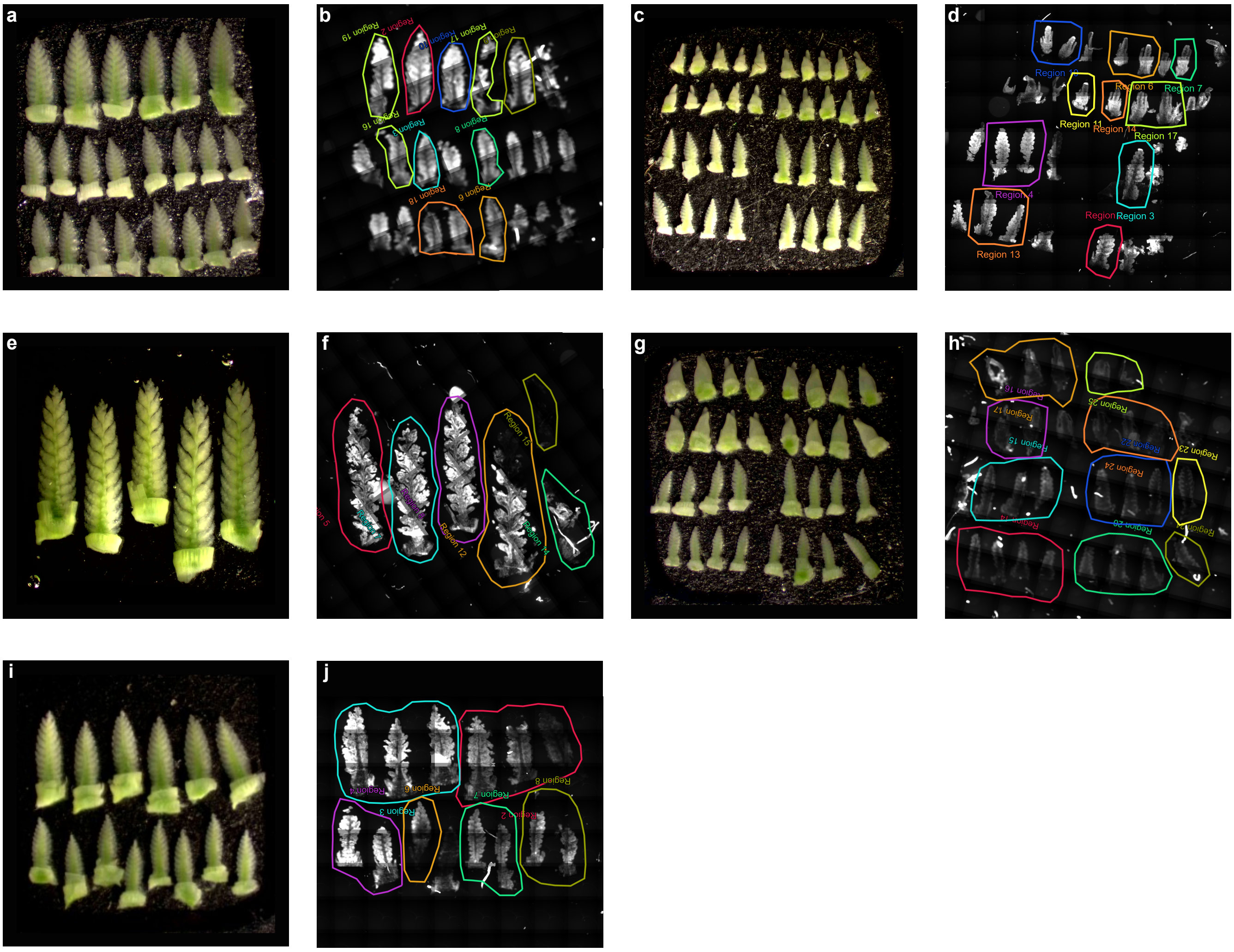


**Supplementary Figure 1. OCT block layout and 'Region of Interest' selections for five MERSCOPE experimental runs.**

Layout of wheat inflorescence in OCT blocks (**a,c,e,g,i**), annotated with genotype and developmental time point annotations. Asterisk annotations denote spikes selected for final analysis. Images of OCT blocks taken on Leica Dissection Microscope. DAPI stain overview and experimental region selections from MERSCOPE Instrument output (**b,d,f,h,j**).

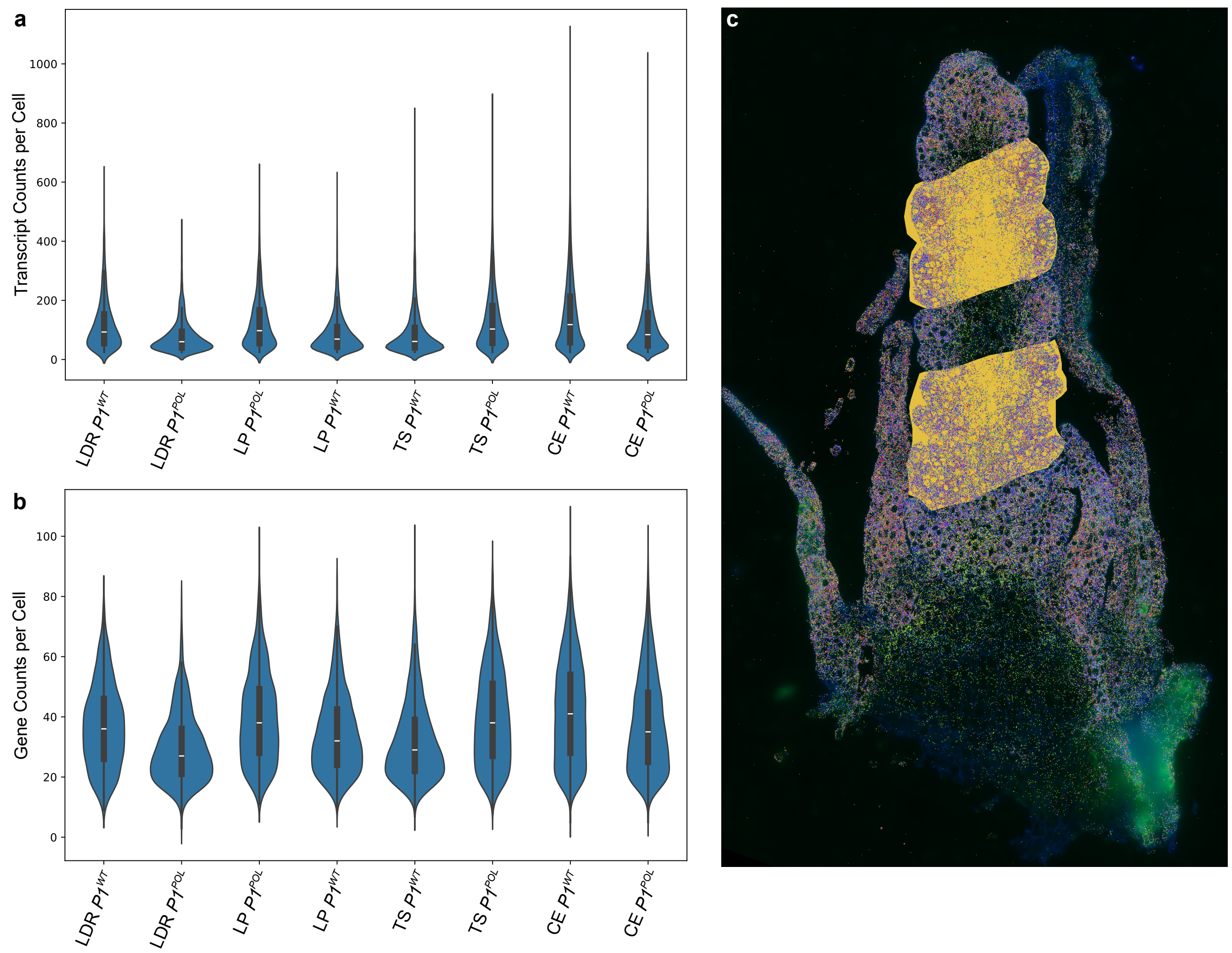


**Supplementary Figure 2. Quality control checks of eight samples and high correlations between MERFISH data and bulk RNA-seq data indicate high sample quality.**

**a,b,** Quality control metrics of (**a**) Total Transcript Counts per cell and (**b**) Total Gene Counts per cell in high quality cells only, in eight samples, across four time points and two genotypes used for onward analysis. **c,** Example of *in-silico* microdissection used to select transcript counts in $P1^{WT}$ LDR (W2.5) inflorescence correlated to bulk RNA-seq data (Supplementary Table 7).

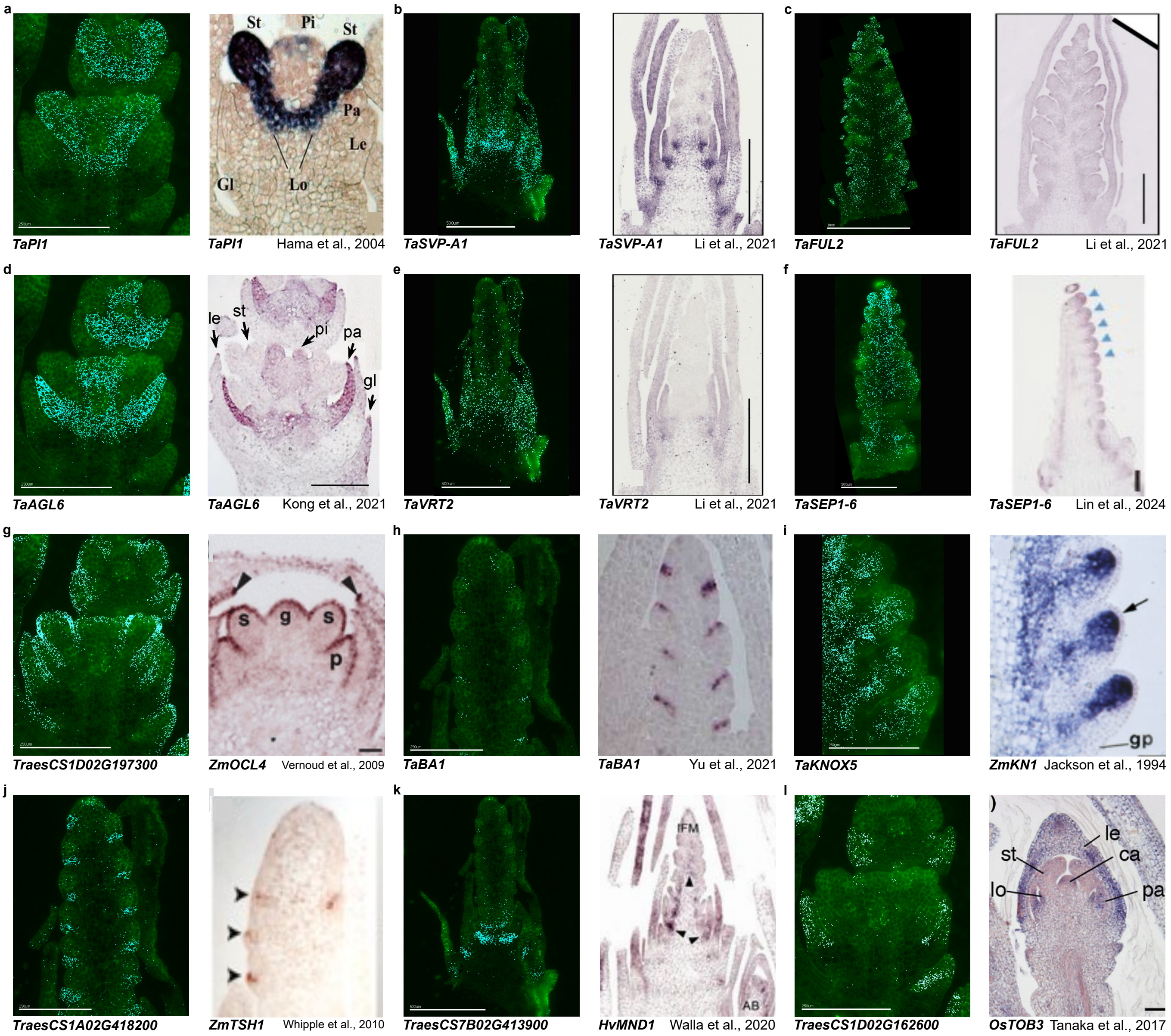


**Supplementary Figure 3. *in-situ* hybridisation results in cereals from equivalent tissues and time points as those used for MERFISH.**

**a-l,** Transcript of wheat genes in MERFISH $P1^{WT}$ samples (left) compared to *in-situ* hybridisation of wheat gene or cereal ortholog at equivalent inflorescence stage in published studies (right). Details of growth stages, gene IDs and publication are found in Supplementary Table 8.

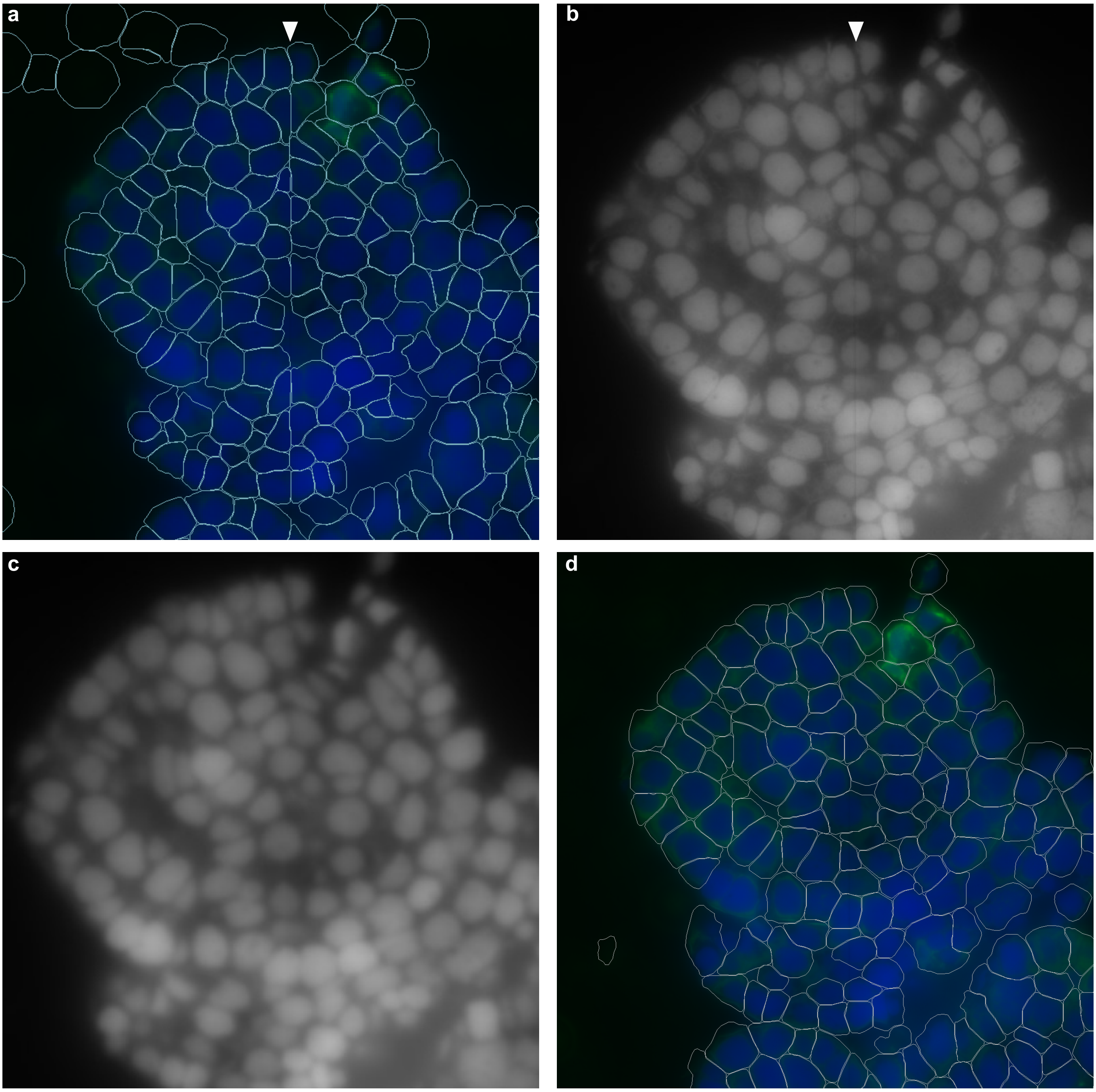


**Supplementary Figure 4. Editing seam lines in staining images improves segmentation along cell boundaries.**

**a,** Segmentation outputs without staining image edits, visualised in Vizgen MERSCOPE Visualiser Tool. DAPI stain in blue, PolyT in green, and cell segmentation boundaries in white. White arrow denotes seam line detected through segmentation resulting in false cell boundaries. **b,** Raw DAPI stain, visualised in ImageJ. White arrow denotes seam line detected through segmentation resulting in false cell boundaries. **c,** Image J filters (Maximum Filter, x3 and Median Filter, x3) applied to stitching edge and Gaussian blur applied to full DAPI stain image. **d,** Cell segmentation of edited staining images, after filtering of high-quality cells.
