## Supplementary Notes for "Spatial Transcriptomics Reveals Expression Gradients in Developing Wheat Inflorescences at Cellular Resolution"

### Supplementary Note 1

Given the hybridisation-based nature of MERFISH, we anticipated assessing hybridisation across highly similar sequences given that wheat's polyploid genome leads to ~ 98% sequence identity among homoeologous genes. Thus, the probes designed for one homoeolog would not discriminate against the other copies. To evaluate this, we included a probe set for a homoeolog triad with high expression levels in the A and D genomes, but no expression of the B genome copy (*TraesCS1B02G448400*). Using a probe set designed against the B genome transcript (96.6% and 97.2% sequence identity against A and D homoeologs, respectively), we detected an average of 0.21 transcripts per cell, eight times higher than the average detection of blank probe sets in this sample, suggesting that the probe effectively detected transcripts from homoeologous copies.

### Supplementary Note 2

The ABCDE model, and the subsequent floral quartet model, have been well established as a core explanation for floral organ development in *Arabidopsis thaliana*[1-3]. Briefly, different flower parts are specified by combinatorial tetrameric protein complexes that activate or repress target genes controlling floral organ development and identity.

The model divides the flower into four whorls containing different organs. The first whorl of floral organs, sepals, is postulated to be specified by A- and E-class protein complexes. The petals form the second whorl and are specified by A-, B-, and E-class proteins[4]. The third whorl, stamens, is determined by B-, C-, and E-class proteins, while the carpel (fourth whorl) is specified by C- and E-class protein complexes[4,5]. The ovule within the carpel is specified by D- and E-class proteins but is not considered a separate whorl[6,7].

In *Arabidopsis thaliana*, the A-class function is encoded by the *APETALA1* (*AP1*) and *APETALA2* (*AP2*) genes; the latter being the only gene in the ABCDE model not belonging to the MIKC$^{c}$-type MADS-box family[8-10]. The B-class function is encoded by the *APETALA3* (*AP3*) and *PISTILLATA* (*PI*) genes[11,12]. The C- and D- class functions are encoded by *AGAMOUS* (*AG*) and several *AGAMOUS*-like genes, namely *SEEDSTICK* (*STK*, also known as *AGL11*), *SHATTERPROOF1* (*SHP1*, also known as *AGL1*) and *SHP2* (*AGL5*)[13,14]. The E-class function is also encoded by several *AGAMOUS*-like genes, namely *SEPALLATA1* (*SEP1*, also known as *AGL2*), *SEP2* (*AGL4*), *SEP3* (*AGL9*), and *SEP4* (*AGL3*)[15,16].

The ABCDE model can also be applied to monocots[17,18]. The floret, a reduced flower typical for grasses, is formed in the axil of a subtending bract (lemma) and consists of an outer perianth (palea), an inner perianth (lodicules), as well as stamens and a gynoecium containing a single ovule[19]. The B-, C-, and D-class genes and functions are highly conserved between mono- and dicots, while there are some differences in A- and E-class genes, which will be discussed below.

### Background

#### B-class

Loss-of-function mutants of the *AP3* orthologs *Superwoman1* (*SPW1*) in rice or *silky1* (*si1*) in maize lead to homeotic conversion of lodicules to palea-like organs and stamens into carpel-like organs[20,21]. Similar phenotypes were observed in the loss-of-function mutant *sterile tassel silky ear1* (*sts1*) in maize and RNAi knockdown of *OsMADS2* and *OsMADS4* in rice, all of which are orthologs of *PI* [22,23].

In barley and wheat, there are two *AP3* orthologs on chromosome groups 6 and 7. In wheat, only the group 7 genes are expressed in whole-tissue RNA-seq datasets, while no expression can be detected for the group 6 genes (expVIP[24,25]; Paragon-*P1*$^{WT}$ NIL RNA-seq data in this manuscript).

#### C-class

Two C-class genes have been identified in rice, namely *OsMADS3* and *OsMADS58*. While loss-of-function mutants of the former gene lead to partial or complete homeotic transformation of stamens into lodicules, the *osmads58* mutant showed no floral defects[26-28]. The double mutant of *osmads3* and *osmads58*, however, has lost floral meristem determinacy and continuously formed lodicules and carpel-like organs instead of stamens and the carpel[26,27].

#### D-class

There are two D-class genes characterized in rice, designated as *OsMADS13* and *OsMADS21*. The *osmads13* loss-of-function mutant shows complete homeotic transformation of the ovule into carpelloid tissue[29]. In contrast, loss-of-function of *osmads21* has no apparent phenotype on rice flowers and the *osmads13 osmads21* double mutant is identical to the *osmads13* mutant, with no additive phenotype observed[27,29]. This suggests that *OsMADS21* has lost its function to regulate ovule identity. In wheat, the orthologs of *OsMADS21* are not expressed during early spike development (Paragon-*P1*$^{WT}$ NIL RNA-seq data in this manuscript). Instead, the genes are expressed, in both wheat and rice, in spike tissue during spike emergence and anthesis as well as in developing grains, suggesting neofunctionalization of this *AG*-paralog (expVIP[24,25])[30-32].

#### E-class

The known genes encoding E-class function in monocots can be divided into three groups: *SEP3*, *LOFSEP*, and *AGL6*-like[33-35].

The two *SEP3* orthologs in rice, *OsMADS7* and *OsMADS8*, have been postulated to act redundantly as knockdown lines of either gene show no or only mild floral phenotypes (carpels in the *osmads7* RNAi line developed three stigmas rather than two)[36]. Similarly, loss-of-function mutants of the *SEP3* ortholog *HvMADS8* in barley have no discernible phenotype under low temperatures, but lose floral determinacy under high temperatures, resulting in the formation of extra carpel-like organs that lack ovules[37]. Simultaneous knockdown of both *OsMADS7* and *OsMADS8*, however, led to homeotic transformation of lodicules into lemma/palea-like structures, elongation of anthers and their filaments, conversion of carpels into green carpel-like organs with trichomes, as well as loss of floral determinacy[36].

The *LOFSEP* clade in rice consists of three genes, namely *OsMADS1*, *OsMADS5*, and *OsMADS34*. The loss-of-function of either *osmads5* or *osmads34* did not alter floral morphology; the only observable phenotype of *osmads34* was the conversion of the sterile lemmas of the rice flower into elongated lemma/leaf-like structures[34,38,39]. By contrast, *osmads1* loss-of-function mutants led to homeotic transformation of lemma, palea, and lodicules into leaf-like organs, as well as either changes to the number of stamens and pistils, conversion of stamens and pistils into leaf-like organs, or loss of floral determinacy leading to formation of multiple flowers or even spikelets[34,38]. Double knockout combinations of *osmads1* with either *osmads5* or *osmads34* showed similar phenotypes to the *osmads1* single mutant, whereas most floral organs in the triple *osmads1 osmads5 osmads34* mutant were transformed into leaf-like structures in addition to the previously mentioned defects of the single *osmads1* mutant[34]. In stark contrast to the results obtained in rice, the triple knockout of the *LOFSEP* genes in barley (*hvmads1 hvmads5 hvmads34*) only resulted in the transformation of lemmas into more leaf-like organs[40].

An *AGL6*-like gene in rice, *OsMADS6*, has been shown to also contribute E-class function, as the loss-of-function mutant led to altered palea morphology, homeotic conversion of lodicules and stamens into glume-like organs, and a loss of floral determinacy[41]. These phenotypes are mimicked in the barley *hvmads6* and durum wheat *Ttagl6* loss-of-function mutants, which also show transformation of palea into lemma-like organs, absence of lodicules or their transformation into lemma-like organs, conversion of two stamens into lemma-like organs in barley, while stamens in wheat carried two anthers, as well as a loss of floral determinacy in both grasses[40,42,43].

### A-class

The A-function of the ABCDE model has been recognized to be more complex than oftentimes portrayed. The classic Arabidopsis *ap1* and *ap2* mutants, which are postulated to control sepal and petal identity, can also cause a variety of other fates for first and second whorl organs, such as complete absence, transformation into bract-like structures, or replacement by secondary flowers[44]. These aspects cannot be explained by the ABCDE model, as they are more likely a result of incomplete transition to or establishment of a floral meristem in the outer whorls, rather than defects in floral organ identity[45].

These limitations of the ABCDE model explain why it has been difficult to identify genes with a "classic" A-function in other species. For monocots, the rice *AP1/FUL*-like genes *OsMADS14* and *OsMADS15* have been proposed to fulfil an A-function in a sense, albeit with some caveats[46]. The single loss-of-function mutants of *osmads14* and *osmads15* showed no strong floral phenotypes, except for a reduction in the size of paleae (without affecting palea identity) and an increase in the length of the sterile lemmas in *osmads15*. The inflorescence of the homozygous *osmads14 osmads15* double mutant aborted development after transitioning to an inflorescence meristem with primary branch primordia. Plants with one functional copy of either gene (*osmads14*;*osmads15*/+ or *osmads14*/+;*osmads15*) showed floral organ identity defects consistent with a loss / reduction of A-function, such as homeotic transformation of paleae into carpel-like organs and lodicules into stamen-like organs[46]. There are two more AP1/FUL-like genes in rice, *OsMADS18* and *OsMADS20*, but their function seems to have diverged as they show low expression levels in floral organs[46].

In wheat, the *AP1/FUL*-like genes *VRN1*, *FUL2*, and *FUL3* have been shown to have a similar function as *OsMADS14* and *OsMADS15* have in rice[47]. The single loss-of-function mutants showed no observable phenotypes in spikes or flowers. The *vrn1 ful2* double and *vrn1 ful2 ful3* triple mutants formed leafy shoots in place of spikelets. In the *vrn1 ful2* double mutant, after forming a "leafy" first floret all subsequent florets were replaced with leaves and finally inflorescence tillers. The floral organs of the first floret were malformed, with leafy paleae and lodicules, reduced number of anthers, anthers fused to ovaries, and multiple ovaries[47]. The *vrn1 ful2 ful3* triple mutant did not form any floral organs, but only produced leafy tillers. Analysis of different mutant combinations showed that as long as one functional allele of *FUL2* was present (e.g. *vrn1*-null/*ful-A2*-null *ful-B2*/+) fertile florets were formed that gave rise to viable grain, suggesting a central role for *FUL2* in spikelet development[47].

Similar difficulties for ascertaining A-function also occur for *AP2* orthologs in wheat. A double knockout of *AP2-like 2* (*AP2L2*) and *AP2L5* in wheat led to transformation of florets into sterile bracts[48]. The most distal florets on a given spikelet did form flowers, but these were malformed and did not produce grain. The observed phenotypes included absence of lodicules and a reduced number of stamens, while some but not all flowers were missing the palea, formed carpel-like organs in place of the lodicules, or produced only carpel-like structures instead of other floral organs. Expression analysis showed that C-class *AG* genes were upregulated in the *ap2l2 ap2l5* double mutant[48]. This is consistent with *AP2* A-function in Arabidopsis, which represses C-class function in the outer whorls[44].

### Spatial Transcriptomics results and interpretation

Building on all these scientific accomplishments, the spatial transcriptomics data presented here allows us to examine the expression of the (proposed) members of the ABCDE model in wheat (Extended Data Fig. 5) and ascertain whether particular genes are predictive of floral organ identity.

As mentioned above, the B-, C-, and D-class genes and their function seem to be well conserved across species. Concordantly, the expression domains of the orthologous genes in wheat overlap with their proposed function. The wheat *AP3* ortholog (orthologous to *OsMADS16*) is expressed in cells of both lodicule and stamen primordia, as are the two *PI* orthologs *PI-1* and *PI-2* (Extended Data Fig. 5a). Likewise, expression of the two C-class genes *AG-1* and *AG-2* (orthologous to *OsMADS2* and *OsMADS4*) is restricted to the stamen and carpel primordia (Extended Data Fig. 5b). Only one of the two D-class genes was included in the MERFISH panel, but the expression of *STK1* (orthologous to *OsMADS13*) can only be detected in carpel primordia (Extended Data Fig. 5c).

We noted divergence in the function of E-class genes, in particular members of the *LOFSEP* clade, between species, even among monocots. The expression pattern for five *SEP1* orthologs in wheat (*SEP1-1*, *SEP1-2*, *SEP1-4*, *SEP1-5*, *SEP1-6*) is scattered throughout the spike, but mostly absent from floral organ primordia (Extended Data Fig. 5e). Instead, most of the expression is localized to cell primordia of glumes and lemmas, which is consistent with loss-of-function phenotypes observed in rice and barley. This suggests that the *LOFSEP* clade in wheat (and possibly within most grasses) is not involved in determining floral organ identity, but instead determines bract (glume, lemma) and spike architecture as was proposed by work in barley[40,49,50].

In contrast, the *SEP3* and *AGL6* orthologs in wheat have overlapping, but not fully redundant, expression domains across all floral organs, suggesting that they fulfil the E-class function in determining floral organ identity as proposed by the ABCDE model (Extended Data Fig. 5d).

Similar to the *LOFSEP* clade, the *AP1/FUL*-like orthologs of wheat do not seem to contribute to floral organ identity, as they are ubiquitously expressed (*VRN1* and *FUL3*) or mostly expressed in bract-like tissues (*FUL2*; Extended Data Fig. 5f). While there is some expression of *FUL2* in palea primordia, it is unlikely a key gene determining palea identity as can be seen from the lack of a phenotype in the *ful2*-null mutant[47]. More likely, the *AP1/FUL*-like genes are involved in determining spike architecture by coordinating meristem phase transition.

Two wheat orthologs of *AP2* are present within the MERFISH panel, namely *AP2-L2* and *AP2-L5*. The latter is expressed in most cells, while *AP2-L2* is mostly localized in lemma, palea, and stamen primordia, but also to some extent within the carpel primordia (Extended Data Fig. 5f). Since the *ap2l2*-null mutant did not show any changes in floral organ identity, it either functions redundantly with other genes or it does not contribute to floral organ identity[47].

The cell segmentation and unsupervised clustering yielded 18 expression domains (EDs) from the MERFISH datasets across the developing inflorescence time-course (Extended Data Fig. 5g). Leaves and bracts are defined by ED1+2, while floral tissues are defined by ED15 (palea), ED17 (lodicules), ED10+14 (stamen), and ED16 (carpel). The overlap of these EDs with well-known floral organ identity genes (Extended Data Fig. 5a-d) highlights the validity of the approach used here and shows the potential for correctly tagging and tracking developing tissues through time and space.

To summarize, the current spatial transcriptomic dataset suggests a conserved function of several MIKC$^c$-type MADS-box genes in the determination of floral organ identity based on comparative analysis with data from other species. The expression domains of B-, C-, D-, and some E-class genes agrees with their putative function in floral organ development. Other proposed A- and E-class members are unlikely to contribute to floral organ development *per se* but rather determine the transition between different meristem fates. These conclusions need to be further tested experimentally through, for example, functional analyses of mutants.

### Supplementary Notes References


1 Ma, H. & dePamphilis, C. The ABCs of floral evolution. *Cell* **101**, 5-8 (2000). https://doi.org/10.1016/S0092-8674(00)80618-2

2 Theissen, G., Melzer, R. & Rumpler, F. MADS-domain transcription factors and the floral quartet model of flower development: linking plant development and evolution. *Development* **143**, 3259-3271 (2016). https://doi.org/10.1242/dev.134080

3 Bowman, J. L. & Moyroud, E. Reflections on the ABC model of flower development. *Plant Cell* **36**, 1334-1357 (2024). https://doi.org/10.1093/plcell/koae044

4 Honma, T. & Goto, K. Complexes of MADS-box proteins are sufficient to convert leaves into floral organs. *Nature* **409**, 525-529 (2001). https://doi.org/10.1038/35054083

5 Coen, E. S. & Meyerowitz, E. M. The war of the whorls: genetic interactions controlling flower development. *Nature* **353**, 31-37 (1991). https://doi.org/10.1038/353031a0

6 Colombo, L. *et al*. The petunia MADS box gene *FBP11* determines ovule identity. *Plant Cell* **7**, 1859-1868 (1995). https://doi.org/10.1105/tpc.7.11.1859

7 Becker, A. & Theissen, G. The major clades of MADS-box genes and their role in the development and evolution of flowering plants. *Mol Phylogenet Evol* **29**, 464-489 (2003). https://doi.org/10.1016/s1055-7903(03)00207-0

8 Mandel, M. A., Gustafson-Brown, C., Savidge, B. & Yanofsky, M. F. Molecular characterization of the *Arabidopsis* floral homeotic gene *APETALA1*. *Nature* **360**, 273-277 (1992). https://doi.org/10.1038/360273a0

9 Jofuku, K. D., den Boer, B. G., Van Montagu, M. & Okamuro, J. K. Control of Arabidopsis flower and seed development by the homeotic gene *APETALA2*. *Plant Cell* **6**, 1211-1225 (1994). https://doi.org/10.1105/tpc.6.9.1211

10 Irish, V. F. The flowering of Arabidopsis flower development. *Plant J* **61**, 1014-1028 (2010). https://doi.org/10.1111/j.1365-313X.2009.04065.x

11 Jack, T., Brockman, L. L. & Meyerowitz, E. M. The homeotic gene *APETALA3* of *Arabidopsis thaliana* encodes a MADS box and is expressed in petals and stamens. *Cell* **68**, 683-697 (1992). https://doi.org/10.1016/0092-8674(92)90144-2

12 Goto, K. & Meyerowitz, E. M. Function and regulation of the *Arabidopsis* floral homeotic gene *PISTILLATA*. *Genes Dev* **8**, 1548-1560 (1994). https://doi.org/10.1101/gad.8.13.1548

13 Favaro, R. *et al*. MADS-box protein complexes control carpel and ovule development in Arabidopsis. *Plant Cell* **15**, 2603-2611 (2003). https://doi.org/10.1105/tpc.015123

14 Pinyopich, A. *et al*. Assessing the redundancy of MADS-box genes during carpel and ovule development. *Nature* **424**, 85-88 (2003). https://doi.org/10.1038/nature01741

15 Pelaz, S., Ditta, G. S., Baumann, E., Wisman, E. & Yanofsky, M. F. B and C floral organ identity functions require *SEPALLATA* MADS-box genes. *Nature* **405**, 200-203 (2000). https://doi.org/10.1038/35012103

16 Ditta, G., Pinyopich, A., Robles, P., Pelaz, S. & Yanofsky, M. F. The *SEP4* gene of *Arabidopsis thaliana* functions in floral organ and meristem identity. *Curr Biol* **14**, 1935-1940 (2004). https://doi.org/10.1016/j.cub.2004.10.028

17 Callens, C., Tucker, M. R., Zhang, D. & Wilson, Z. A. Dissecting the role of MADS-box genes in monocot floral development and diversity. *J Exp Bot* **69**, 2435-2459 (2018). https://doi.org/10.1093/jxb/ery086

18 Dreni, L. The ABC of Flower Development in Monocots: The Model of Rice Spikelet. *Methods Mol Biol* **2686**, 59-82 (2023). https://doi.org/10.1007/978-1-0716-3299-4_3

19 Kellogg, E. A. Genetic control of branching patterns in grass inflorescences. *Plant Cell* **34**, 2518-2533 (2022). https://doi.org/10.1093/plcell/koac080

20 Nagasawa, N. *et al*. *SUPERWOMAN1* and *DROOPING LEAF* genes control floral organ identity in rice. *Development* **130**, 705-718 (2003). https://doi.org/10.1242/dev.00294

21 Ambrose, B. A. *et al*. Molecular and genetic analyses of the *silky1* gene reveal conservation in floral organ specification between eudicots and monocots. *Mol Cell* **5**, 569-579 (2000). https://doi.org/10.1016/s1097-2765(00)80450-5

22 Yao, S. G., Ohmori, S., Kimizu, M. & Yoshida, H. Unequal genetic redundancy of rice *PISTILLATA* orthologs, *OsMADS2* and *OsMADS4*, in lodicule and stamen development. *Plant Cell Physiol* **49**, 853-857 (2008). https://doi.org/10.1093/pcp/pcn050

23 Bartlett, M. E. *et al*. The Maize *PI/GLO* Ortholog *Zmm16/sterile tassel silky ear1* Interacts with the Zygomorphy and Sex Determination Pathways in Flower Development. *Plant Cell* **27**, 3081-3098 (2015). https://doi.org/10.1105/tpc.15.00679

24 Ramirez-Gonzalez, R. H. *et al.* The transcriptional landscape of polyploid wheat. *Science* **361** (2018). https://doi.org/10.1126/science.aar6089
25 Borrill, P., Ramirez-Gonzalez, R. & Uauy, C. expVIP: a Customizable RNA-seq Data Analysis and Visualization Platform. *Plant Physiol* **170**, 2172-2186 (2016). https://doi.org/10.1104/pp.15.01667
26 Sugiyama, S. H., Yasui, Y., Ohmori, S., Tanaka, W. & Hirano, H. Y. Rice Flower Development Revisited: Regulation of Carpel Specification and Flower Meristem Determinacy. *Plant Cell Physiol* **60**, 1284-1295 (2019). https://doi.org/10.1093/pcp/pcz020
27 Dreni, L. *et al.* Functional analysis of all *AGAMOUS* subfamily members in rice reveals their roles in reproductive organ identity determination and meristem determinacy. *Plant Cell* **23**, 2850-2863 (2011). https://doi.org/10.1105/tpc.111.087007
28 Yamaguchi, T. *et al.* Functional diversification of the two C-class MADS box genes *OSMADS3* and *OSMADS58* in *Oryza sativa*. *Plant Cell* **18**, 15-28 (2006). https://doi.org/10.1105/tpc.105.037200
29 Dreni, L. *et al.* The D-lineage MADS-box gene *OsMADS13* controls ovule identity in rice. *Plant J* **52**, 690-699 (2007). https://doi.org/10.1111/j.1365-313X.2007.03272.x
30 Wang, H. *et al.* Analysis of non-coding transcriptome in rice and maize uncovers roles of conserved lncRNAs associated with agriculture traits. *Plant J* **84**, 404-416 (2015). https://doi.org/10.1111/tpj.13018
31 Sakai, H. *et al.* Rice Annotation Project Database (RAP-DB): an integrative and interactive database for rice genomics. *Plant Cell Physiol* **54**, e6 (2013). https://doi.org/10.1093/pcp/pcs183
32 Kawahara, Y. *et al.* Improvement of the *Oryza sativa* Nipponbare reference genome using next generation sequence and optical map data. *Rice (N Y)* **6**, 4 (2013). https://doi.org/10.1186/1939-8433-6-4
33 Malcomber, S. T. & Kellogg, E. A. *SEPALLATA* gene diversification: brave new whorls. *Trends Plant Sci* **10**, 427-435 (2005). https://doi.org/10.1016/j.tplants.2005.07.008
34 Wu, D. *et al.* Loss of LOFSEP Transcription Factor Function Converts Spikelet to Leaf-Like Structures in Rice. *Plant Physiol* **176**, 1646-1664 (2018). https://doi.org/10.1104/pp.17.00704
35 Dreni, L. & Ferrandiz, C. Tracing the Evolution of the *SEPALLATA* Subfamily across Angiosperms Associated with Neo- and Sub-Functionalization for Reproductive and Agronomically Relevant Traits. *Plants (Basel)* **11** (2022). https://doi.org/10.3390/plants11212934
36 Cui, R. *et al.* Functional conservation and diversification of class E floral homeotic genes in rice (*Oryza sativa*). *Plant J* **61**, 767-781 (2010). https://doi.org/10.1111/j.1365-313X.2009.04101.x
37 Shen, C. *et al.* *MADS8* is indispensable for female reproductive development at high ambient temperatures in cereal crops. *Plant Cell* **36**, 65-84 (2023). https://doi.org/10.1093/plcell/koad246
38 Gao, X. *et al.* The *SEPALLATA*-like gene *OsMADS34* is required for rice inflorescence and spikelet development. *Plant Physiol* **153**, 728-740 (2010). https://doi.org/10.1104/pp.110.156711
39 Hu, Y. *et al.* Interactions of *OsMADS1* with Floral Homeotic Genes in Rice Flower Development. *Mol Plant* **8**, 1366-1384 (2015). https://doi.org/10.1016/j.molp.2015.04.009
40 Shen, C., Yang, X., Wang, D., Li, G. & Tucker, M. R. Functional retrogression of LOFSEPs in specifying floral organs in barley. *aBIOTECH* (2024). https://doi.org/10.1007/s42994-024-00182-4
41 Li, H. *et al.* The *AGL6*-like gene *OsMADS6* regulates floral organ and meristem identities in rice. *Cell Res* **20**, 299-313 (2010). https://doi.org/10.1038/cr.2009.143
42 Sun, M. *et al.* A novel type of malformed floral organs mutant in barley was conferred by loss-of-function mutations of the MADS-box gene *HvAGL6*. *Plant J* **119**, 2609-2621 (2024). https://doi.org/10.1111/tpj.16936
43 Kong, X. *et al.* The wheat *AGL6*-like MADS-box gene is a master regulator for floral organ identity and a target for spikelet meristem development manipulation. *Plant Biotechnol J* **20**, 75-88 (2022). https://doi.org/10.1111/pbi.13696
44 Causier, B., Schwarz-Sommer, Z. & Davies, B. Floral organ identity: 20 years of ABCs. *Semin Cell Dev Biol* **21**, 73-79 (2010). https://doi.org/10.1016/j.semcdb.2009.10.005
45 Litt, A. An Evaluation of A-Function: Evidence from the *APETALA1* and *APETALA2* Gene Lineages. *International Journal of Plant Sciences* **168**, 73-91 (2007). https://doi.org/10.1086/509662
46 Wu, F. *et al.* The ABCs of flower development: mutational analysis of *AP1*/*FUL*-like genes in rice provides evidence for a homeotic (A)-function in grasses. *Plant J* **89**, 310-324 (2017). https://doi.org/10.1111/tpj.13386
47 Li, C. *et al.* Wheat *VRN1*, *FUL2* and *FUL3* play critical and redundant roles in spikelet development and spike determinacy. *Development* **146** (2019). https://doi.org/10.1242/dev.175398

48 Debernardi, J. M., Greenwood, J. R., Jean Finnegan, E., Jernstedt, J. & Dubcovsky, J. *APETALA 2*-like genes *AP2L2* and *Q* specify lemma identity and axillary floral meristem development in wheat. *Plant J* **101**, 171-187 (2020). https://doi.org/10.1111/tpj.14528
49 Zhang, Y. *et al. MADS1*-regulated lemma and awn development benefits barley yield. *Nat Commun* **15**, 301 (2024). https://doi.org/10.1038/s41467-023-44457-8
50 Li, G. *et al. MADS1* maintains barley spike morphology at high ambient temperatures. *Nat Plants* **7**, 1093-1107 (2021). https://doi.org/10.1038/s41477-021-00957-3
